# YTHDF2 in dentate gyrus is the m^6^A reader mediating m^6^A modification in hippocampus-dependent learning and memory

**DOI:** 10.1101/2022.10.31.514574

**Authors:** Mengru Zhuang, Xiaoqi Geng, Peng Han, Fanghao Liang, Pengfei Che, Chao Liu, Lixin Yang, Jun Yu, Zhuxia Zhang, Wei Dong, Sheng-Jian Ji

## Abstract

*N*^*6*^-methyladenosine (m^6^A) has been demonstrated to regulate learning and memory in mice. To investigate the mechanism by which m^6^A modification exerts its function through its reader proteins in the hippocampus, as well as to unveil the specific subregions of the hippocampus that are crucial for memory formation, we generated dentate gyrus (DG)-, CA3-, and CA1-specific *Ythdf1* and *Ythdf2* conditional knockout (cKO) mice, respectively. Surprisingly, we found that only the DG-specific *Ythdf2* cKO mice displayed impaired memory formation, which is inconsistent with the previous report showing that YTHDF1 was involved in this process. YTHDF2 controls the stability of its target transcripts which encode proteins that regulate the elongation of mossy fibers (MF), the axons of granule cells in DG. DG-specific *Ythdf2* ablation caused MF overgrowth and impairment of the MF-CA3 excitatory synapse development and transmission in the stratum lucidum. Thus, this study identifies the m^6^A reader YTHDF2 in dentate gyrus as the only regulator that mediates m^6^A modification in hippocampus-dependent learning and memory.

## Introduction

The hippocampus is crucial for encoding and storing diverse memories during the episodic events ^1-3^. Hippocampus receives input from and forms connections with the entorhinal cortex (EC), and is mainly composed of dentate gyrus (DG) and cornu ammonis (CA3, CA2, and CA1) subregions ^4^. In the classical model of corticohipocampal circuits, neurons in the superficial entorhinal cortex send glutamatergic projections by perforant path (PP) to the granule cells of DG, whose mossy fibers (MF) excite CA3 neurons and the excited CA3 neurons in turn passage the signal to CA1 through Schaffer collateral (SC) which sends the signal back to EC via subiculum ^5^. However, how each of these hippocampal subregions contributes to the intra-hippocampal DG-CA3-CA1 trisynaptic circuit is largely unknown.

*N*6-methyladenosine (m^6^A) is the most abundant RNA modification and regulates RNA fate in the epitranscriptome level ^6^. m^6^A modification has been shown to regulate nervous system development and function ^7,8^. Specifically, m^6^A modification has been reported to regulate memory formation and consolidation in mouse models ^9-12^. Proper methylation of transcripts in mouse hippocampus is critical for memory consolidation since ablation of the m^6^A writer METTL13 led to a reduction in the memory consolidation capacity ^11^. The fate of m^6^A-modified transcripts is mainly decoded by their readers, thus identification and characterization of the specific m^6^A readers in the hippocampus are necessary to elucidate the involvement of m^6^A modification during memory storage and retrieval.

The m^6^A reader YTHDF1 has been shown to mediate the m^6^A function in learning and memory, which was demonstrated by impaired long-term potentiation and synaptic transmission in the *Ythdf1* null mice ^12^. However, the potential side-effects brought by disfunctions of other brain areas besides the hippocampus or other areas outside of the brain when using the null mutant mouse model should be minded, given that encoding and retrieving of spatial memories rely on the neural circuits formed between hippocampal and cortical areas of the brain ^4,13^. Besides, the role of another major m^6^A reader YTHDF2 in regulating hippocampus-based memory consolidation remains elusive since the *Ythdf2* null mutant mouse is embryonically lethal ^14^. In addition, which component(s) of the DG-CA3-CA1 trisynaptic circuit in the hippocampus is the major player in this process is also unaddressed. Thus, conditional knockout mice in which the genetic ablation of the m^6^A readers is restricted to specific hippocampal subregions are favored to answer these questions.

In this study, we generated DG-, CA3-, and CA1-specific *Ythdf1* and *Ythdf2* knockout mice, respectively. In contrast to the previous report, we surprisingly found that conditional knockout of *Ythdf1* from DG, CA3, or CA1 subregions in the hippocampus did not impair learning and memory. Only DG-specific but not CA3- or CA1-specific *Ythdf2* knockout mice showed defects in memory consolidation. Loss-of-function of YTHDF2 in DG granule cells caused overgrowth of mossy fibers and defects of excitatory DG-CA3 synapses in DG-specific *Ythdf2* knockout hippocampus. The potential target transcripts through which YTHDF2 regulates DG axon growth were identified and verified. Our study identifies YTHDF2 in DG as the only responsible m^6^A reader which mediates the function of m^6^A modification in learning and memory.

## Results

### YTHDF1 and YTHDF2 are highly expressed in the early postnatal mouse hippocampus

The most abundance of m^6^A modification was reported in the brain ^15^. Within the mouse brain, we found that m^6^A has exceptionally high enrichment in the hippocampus, which was demonstrated by the strong immunofluorescence (IF) signals in granule cells in DG and pyramidal cells in CA areas of postnatal day 1 to 50 (P1∼P50) mouse hippocampi using an m^6^A-specific antibody (Supplementary Fig. 1a). The relative m^6^A levels in the P1-P50 mouse hippocampi were further quantified by anti-m^6^A dot blots (Supplementary Fig. 1b, c). We further checked the expression of the m^6^A readers in the P1-P50 mouse hippocampi and their subregions. The mRNA levels of *Ythdfs* in these subregions are varied, which were shown by our analysis (Supplementary Fig. 1d) and the previous reports ^16,17^. Using immunofluorescence, we found that YTHDF1 and YTHDF2 are highly expressed in the mature DG granule cells and all CA pyramidal cells of P1∼P50 mouse hippocampi (Fig. 1a, b). We further carried out Western Blotting of YTHDF1 and YTHDF2 in P1∼P50 mouse hippocampi, which showed the highest expression in the early postnatal hippocampus and the gradual decrease toward adulthood (Fig. 1c-e). The abundant m^6^A and highly expressed reader proteins in the early postnatal hippocampus suggest that m^6^A modification might be involved in the postnatal hippocampal development and functions.

**Fig. 1.**
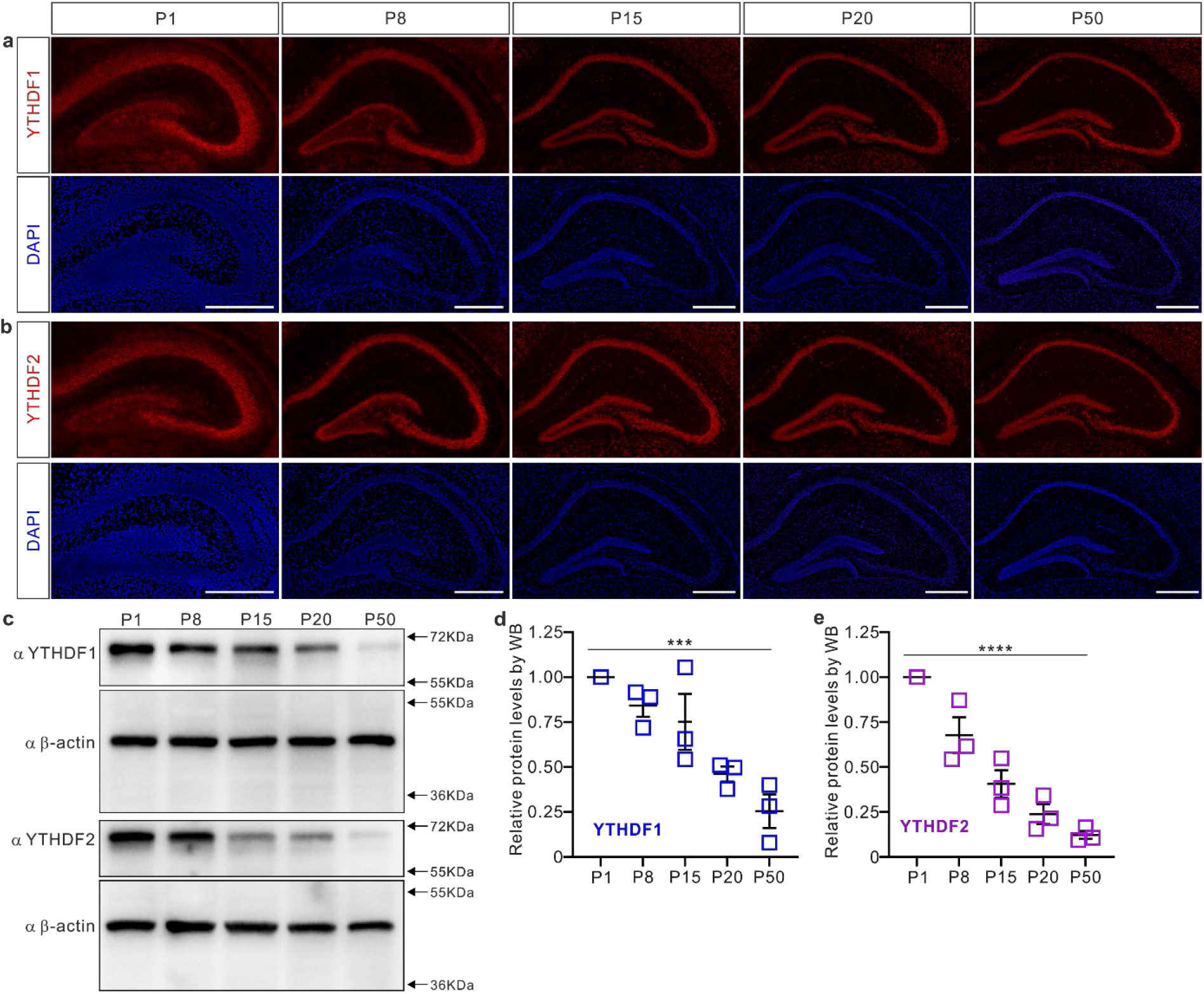
YTHDF1 and YTHDF2 are highly expressed in the early postnatal mouse hippocampus. **a, b**, Representative immunofluorescent images showed that YTHDF1 (a) and YTHDF2 (b) are expressed in the mature DG granule cells and all CA pyramidal cells in P1, P8, P15, P20, and P50 mouse hippocampi. **c-e**, WB shows that expression levels of YTHDF1 and YTHDF2 are highest in the early postnatal hippocampus and gradually decrease in adulthood (c). Quantification of WB is shown in dot blots (d, e): *n* = 3 mice for each stage, \*\*\**P* = 8.54E-4 in d, \*\*\*\**P* = 1.03E-5 in e, by one-way ANOVA. Scale bars, 500 μm (a, b).

### *Ythdf1* and *Ythdf2* are specifically ablated from hippocampal subregions using DG-, CA3-, and CA1-specific Cre lines, respectively

To test whether and how the m^6^A readers YTHDF1 and YTHDF2 work in specific hippocampal subregions to regulate hippocampus-dependent learning and memory, we sought to generate subregion-specific conditional knockouts (cKO) of these proteins. For this purpose, we chose *Pomc-cre* ^18^, *Grik4-cre* ^19^, and *Camk2α-cre* (T29-1) ^20^ lines, which have been used by multiple studies to do DG-, CA3-, or CA1-specific analysis, respectively ^18-26^. By crossing with the *Rosa26-eYFP* reporter ^27^, we validated the hippocampal subregion-specific expression of these Cre lines (Supplementary Fig. 2). Using these Cre lines, we generated DG-, CA3-, and CA1-specific *Ythdf1* or *Ythdf2* conditional knockout mice, respectively (Supplementary Fig. 3). These six cKO mice are viable, fertile, and develop normally with body weights, brain sizes, and morphology comparable to their littermate controls (*Ythdf1*^*fl/fl*^ or *Ythdf2*^*fl/fl*^) (Supplementary Fig. 4). Thus, these hippocampal subregion-specific cKO mice can be used in various behavioral tests to explore the roles and mechanisms of YTHDF1 and YTHDF2 in regulating brain functions.

### Only *Ythdf2* conditional knockout in dentate gyrus but not in other subregions of hippocampus or *Ythdf1* conditional knockouts impairs learning and memory

With these *Ythdf1* and *Ythdf2* cKO mice, we first tested their exploratory behavior, short-term memory, anxiety level, and motor coordination using the open field, novel object recognition, elevated plus-maze, and rotarod tests, respectively. In these tests, the performance of these cKO mice was essentially normal when compared with their control littermates (Supplementary Fig. 5 and 6).

Then the Morris water maze test was adopted to examine the hippocampus-dependent spatial memory of these cKO mice lines. Surprisingly, we found that knockout of *Ythdf1* from either of the three hippocampal subregions caused no defect in memory (Supplementary Fig. 7), suggesting that YTHDF1 is dispensable for hippocampus-dependent learning and memory. For *Ythdf2* cKO, we found that the mice with specific ablation of *Ythdf2* from DG took more time to find the hidden platform in the first three training days and spent significantly less amount of time in the target quadrant in the probe test I on day 3 than their control peers, indicating that they failed to remember the location of the platform (Fig. 2a, b). These DG-specific *Ythdf2* cKO mice learned gradually and achieved comparable performance during training days 4 and 5 as well as in the probe test II on day 6 (Fig. 2a, c). In contrast, CA3- or CA1-specific *Ythdf2* cKO mice showed no defect in the Morris water maze test (Fig. 2d-i).

**Fig. 2.**
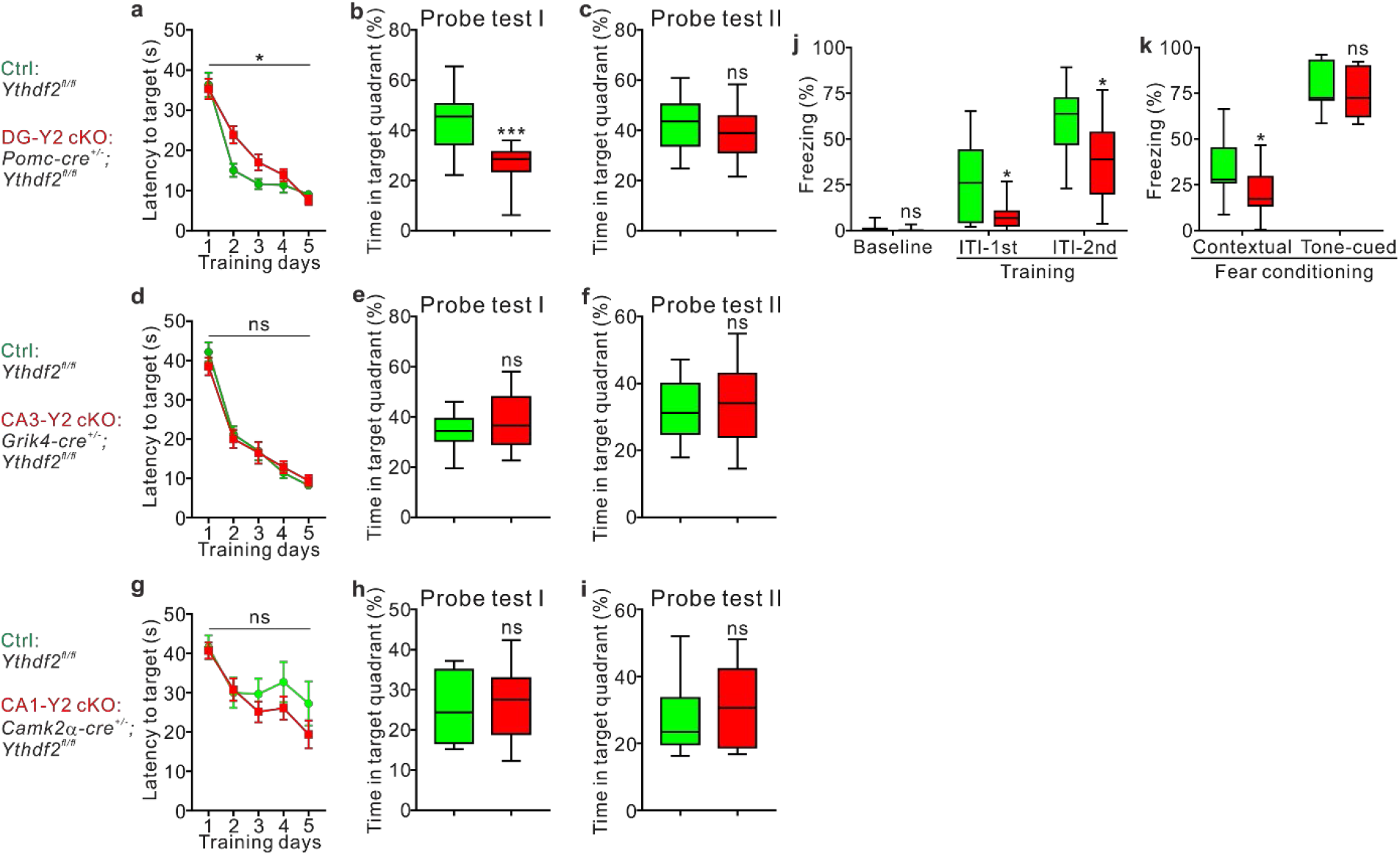
*Ythdf2* conditional knockout in dentate gyrus but not in other hippocampal subregions disturbs learning and memory. DG-, CA3-, and CA1-specific *Ythdf2* cKO are designated as DG-Y2 cKO (a-c, j-k), CA3-Y2 cKO (d-f), and CA1-Y2 cKO (g-i). **a-i**, *Ythdf2* cKO in dentate gyrus but not in other hippocampal subregions impairs learning and memory in the Morris water maze test. The DG-Y2 cKO mice had impaired memory recalling during training days 2 and 3 (a), spent less time in the target quadrant at Probe test I on day 3 (b), and resumed to the comparable level with control (Ctrl) mice at Probe test II on day 6 (c). Both CA3-Y2 and CA1-Y2 cKO mice performed normally in the Morris water maze test (d-f, g-i). **j, k**, *Ythdf2* cKO in dentate gyrus impairs the contextual fear conditioning memory but not the tone-cued fear conditioning memory. The DG-Y2 cKO mice showed less freezing time in the first and second intertrial intervals (ITI) on the training day and when exposed to the contextual conditioning, but behaved normally under the tone-cued conditioning, compared with the control mice. Quantifications of the learning curves of the Morris water maze test are shown in dot plots (mean ± SEM), while the Probe tests I-II of the Morris water maze test and the fear conditioning test are represented as box and whisker plots. For DG-Y2 cKO, *n* = 14 mice for Ctrl and *n* = 10 mice for DG-Y2 cKO in a-c, \**P* = 0.021 in a, \*\*\**P* = 3.23E-4 in b; for CA3-Y2 cKO, *n* = 16 mice for Ctrl and *n* = 14 mice for CA3-Y2 cKO in d-f; for CA1-Y2 cKO, *n* = 12 mice for Ctrl and *n* = 13 mice for CA1-Y2 cKO in g-i; *n* = 11 mice for Ctrl and *n* = 15 mice for DG-Y2 cKO in j and k, \**P* = 0.035 for ITI-1st in j, \**P* = 0.019 for ITI-2nd in j, \**P* = 0.038 for contextual fear conditioning in k; ns, not significant; group difference in dot plots by two-way ANOVA with Tukey’s post hoc test; all box and whisker plots by unpaired Student’s t-test.

To further confirm that hippocampus-dependent learning and memory are indeed disrupted in the DG-specific *Ythdf2* cKO mice, we continued to carry out the fear conditioning. As shown in Fig. 2j, k, the DG-specific *Ythdf2* cKO mice showed less freezing time in the two intertrial intervals (ITIs) and in the contextual fear conditioning test, but not in the tone-cued conditioning test, when compared with the control mice. These results indicate that the contextual fear memory in the cKO mice was impaired, while the tone-cued fear memory in them was not affected, which supports the dysfunction of the hippocampus in the cKO mice because the hippocampus has been believed to be involved in the formation of contextual conditioning fear memory but not the cued conditioning memory ^28^.

In summary, these data suggest that only YTHDF2 in DG but not in other hippocampal subregions or YTHDF1 is required for hippocampus-dependent learning and memory.

### DG-specific *Ythdf2* knockout causes overgrowth of mossy fiber and impairs the development and transmission of DG-CA3 synapses

To understand the mechanism underlying the impaired hippocampus-dependent learning and memory in DG-specific *Ythdf2* knockout mice, we first analyzed whether *Ythdf2* ablation in DG affected early postnatal development of the hippocampus. No obvious defect was detected in the differentiation, neurogenesis, formation, or maturation of DG in the DG-specific *Ythdf2* cKO hippocampus (Supplementary Fig. 8a-h).

We next asked whether the neurite development of GCs was affected by *Ythdf2* ablation. We used Golgi staining to examine the dendrite branching and dendritic spines of GCs, which showed no significant difference between DG-specific *Ythdf2* cKO and control mice (Supplementary Fig. 8i-n). We continued to test whether YTHDF2 regulated GC axon growth. We generated lentiviral shRNAs against *Ythdf2*, which could efficiently knock down YTHDF2 (Fig. 3a, b). Knockdown of YTHDF2 significantly enhanced GC axon growth (Fig. 3c, d). We continued to culture DG-specific *Ythdf2* cKO GC neurons which showed enhanced axon growth compared with control neurons (Fig. 3e). This phenotype was rescued by overexpression of YTHDF2 using a lentiviral overexpression system (Fig. 3e), suggesting the direct regulation of YTHDF2 in GC axon growth. To verify this result in vivo, we used the reporter *Rosa26mT/mG* ^29^ to label mossy fibers. Compared with control mice, DG-specific *Ythdf2* cKO mice showed enhanced growth of mossy fibers (MF): at P5, the length of MF trajectory and the number of GC axons projecting into the CA3 area were remarkably increased (Fig. 3f-h); at P8, the relative area and thickness of MF terminals were robustly increased (Fig. 3i-k), which was further verified by analysis using an MF terminal marker SPO (Supplementary Fig. 9a-c). These data suggest that DG-specific *Ythdf2* cKO caused the overgrowth of mossy fibers.

**Fig. 3.**
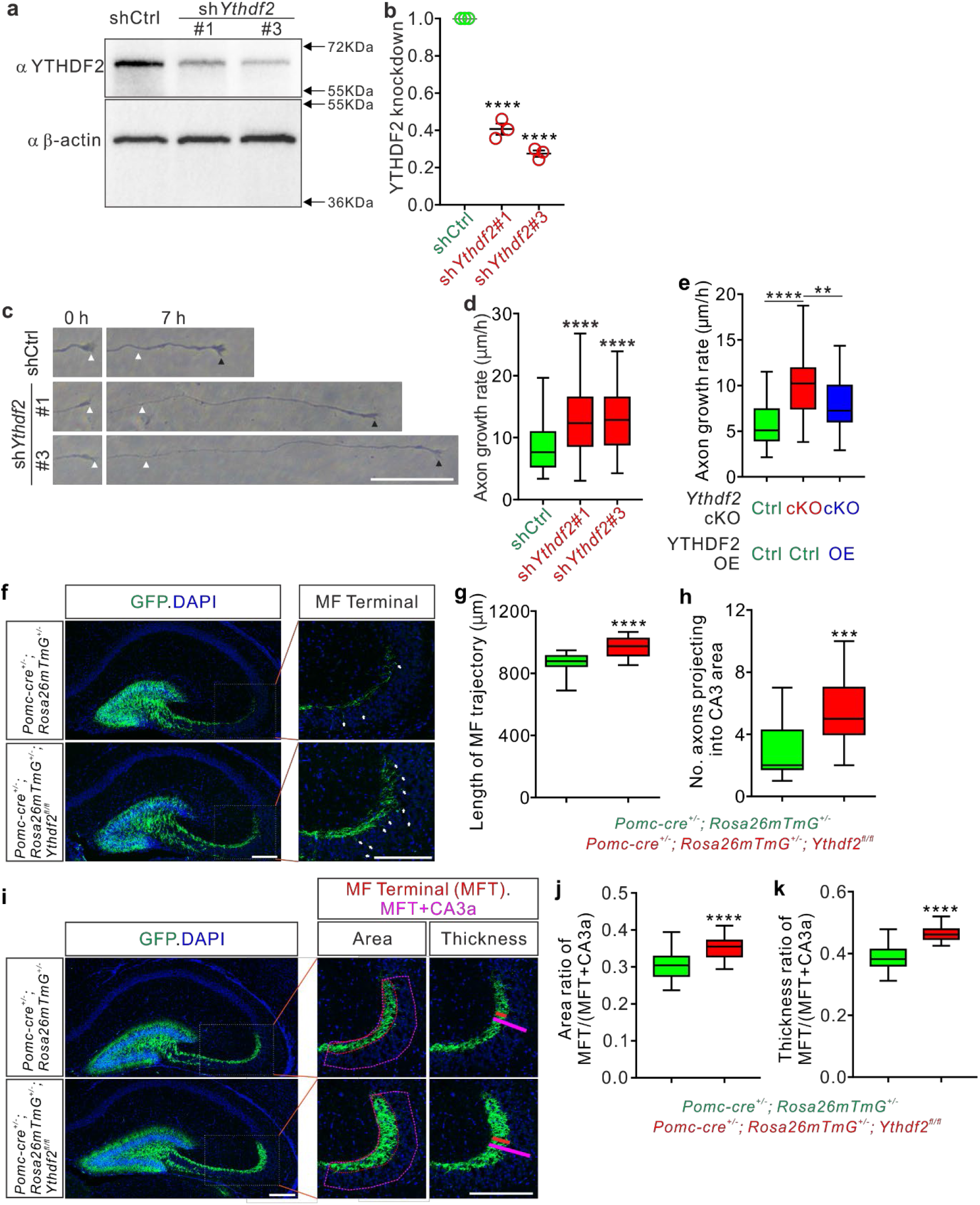
Loss-of-function of YTHDF2 in DG granule cells enhances axon growth both in vitro and in vivo. **a, b**, Knockdown of YTHDF2 in GCs using lentiviral sh*Ythdf2*. Dissociated GCs from wild-type P6-8 mouse hippocampus were infected with lentivirus expressing sh*Ythdf2*#1 and sh*Ythdf2*#3, respectively. Western blotting (a) results showed efficient knockdown (KD) of YTHDF2, indicated by quantification (b). **c, d**, YTHDF2 KD enhanced GC axon growth. GCs cultured in the microfluidic chambers were infected with sh*Ythdf2*. GC axons were imaged at two time points and the axon terminals at 0 and 7 hr were indicated with white and black arrowheads, respectively (c). Axon growth rates were measured (d). **e**, Overgrowth of GC axons in DG-specific *Ythdf2* cKO could be rescued by overexpression of YTHDF2. GCs from control and DG-specific *Ythdf2* cKO mice were cultured in the microfluidic chambers and were infected with lentiviral control or YTHDF2 overexpression construct. GC axons were imaged at two time points and axon growth rates were calculated. **f-h**, DG-specific knockout of *Ythdf2* increased mossy fiber (MF) growth at P5. Representative images showed GFP-labelled MF in control and DG-specific *Ythdf2* cKO mice (f). Arrows pointed to the axons that projected into the CA3 area. Compared with control mice, the length of MF trajectory was significantly increased and more axons were found to project into the CA3 area in the cKO mice. Quantifications were shown in g and h. **i-k**, At P8, the area and thickness of the MF terminal (MFT) in the cKO mice were both increased when compared with control mice. The red and magenta dotted boundaries indicate the MFT area and the “MFT+CA3a” area, respectively, while the red and magenta bars measure the thickness of MFT and “MFT+CA3a”, respectively (i). Quantifications were shown in j and k. All quantifications are represented as box and whisker plots except in b which was dot plot with mean ± SEM. In b, *n* = 3 replicates, shCtrl vs sh*Ythdf2*#1, \*\*\*\**P* = 1.86E-6; shCtrl vs sh*Ythdf2*#3, \*\*\*\**P* = 6.88E-7. In d, shCtrl (*n* = 63 axons) vs sh*Ythdf2*#1 (*n* = 63 axons), \*\*\*\**P* = 1.86E-6; shCtrl vs sh*Ythdf2*#3 (*n* = 62 axons), \*\*\*\**P* = 6.88E-7. In e, *Ythdf2*-cKO/OE-Ctrl (*n* = 49 axons) vs Ctrl/OE-Ctrl (*n* = 27 axons), \*\*\*\**P* = 3.17E-7; *Ythdf2*-cKO/YTHDF2-OE (*n* = 27 axons) vs *Ythdf2*-cKO/OE-Ctrl, \*\**P* = 0.0081. In g and h, *n* = 22 sections for *Pomc-cre*^*+/-*^; *Rosa26mT/mG*^*+/-*^, *n* = 23 sections for *Pomc-cre*^*+/-*^; *Ythdf2*^*fl/fl*^; *Rosa26mT/mG*^*+/-*^, \*\*\*\**P* = 1.73E-6 (g), \*\*\**P* = 1.10E-5 (h). In j and k, *n* = 44 sections for *Pomc-cre*^*+/-*^; *Rosa26mT/mG*^*+/-*^; *n* = 37 sections for *Pomc-cre*^*+/-*^; *Ythdf2*^*fl/fl*^; *Rosa26mT/mG*^*+/-*^, \*\*\*\**P* = 9.06E-17 (j), \*\*\*\**P* = 4.66E-8 (k). b, d, and e by one-way ANOVA followed by Tukey’s post hoc test; all others by unpaired Student’s t-test. At least 3 mice were analyzed for each genotype in each experiment. Scale bars, 50 μm (c), 200 μm (f, i).

We further zoomed in to check the morphology of MF terminals located in the CA3 stratum lucidum, where the excitatory synapses are formed between MF boutons and CA3 dendritic spines. We found that DG-specific *Ythdf2* cKO mice showed a significant reduction in the density of MF boutons compared with their littermate controls (Fig. 4a-c), without changing the MF bouton size (Supplementary Fig. 9d, e). We continued to check the excitatory synapses formed in the stratum lucidum area, which transmit the excitatory signals from DG to CA3. The synapses were identified by co-staining of the presynaptic marker, Bassoon, and the postsynaptic marker, PSD95. As shown in Fig. 4d, e, the numbers of MF-CA3 excitatory synapses in the stratum lucidum were strikingly declined in the cKO mice compared with control, consistent with the decrease of MF bouton density. To further investigate the effects of DG-specific *Ythdf2* cKO on the function of MF-CA3 excitatory synapses, we performed whole-cell voltage clamp recordings in CA3 pyramidal neurons. The miniature excitatory postsynaptic currents (mEPSCs) of CA3 pyramidal neurons were recorded in the presence of 1 μM TTX. As shown in Fig. 4f-h, *Ythdf2* cKO led to decreased frequency of mEPSCs, which further confirms the reduced number of MF-CA3 excitatory synapses in the cKO mice. However, the amplitude of mEPSCs is not changed (Fig. 4i), indicating the postsynaptic receptors are not affected by *Ythdf2* cKO. We also recorded the evoked EPSCs (eEPSCs) triggered by stimulating the DG granule cells. Compared with control synapses, the cKO synapses showed significantly decreased amplitude for eEPSCs (Fig. 4j, k), suggesting that the MF-CA3 synaptic transmission in the *Ythdf2* cKO was impaired.

**Fig. 4.**
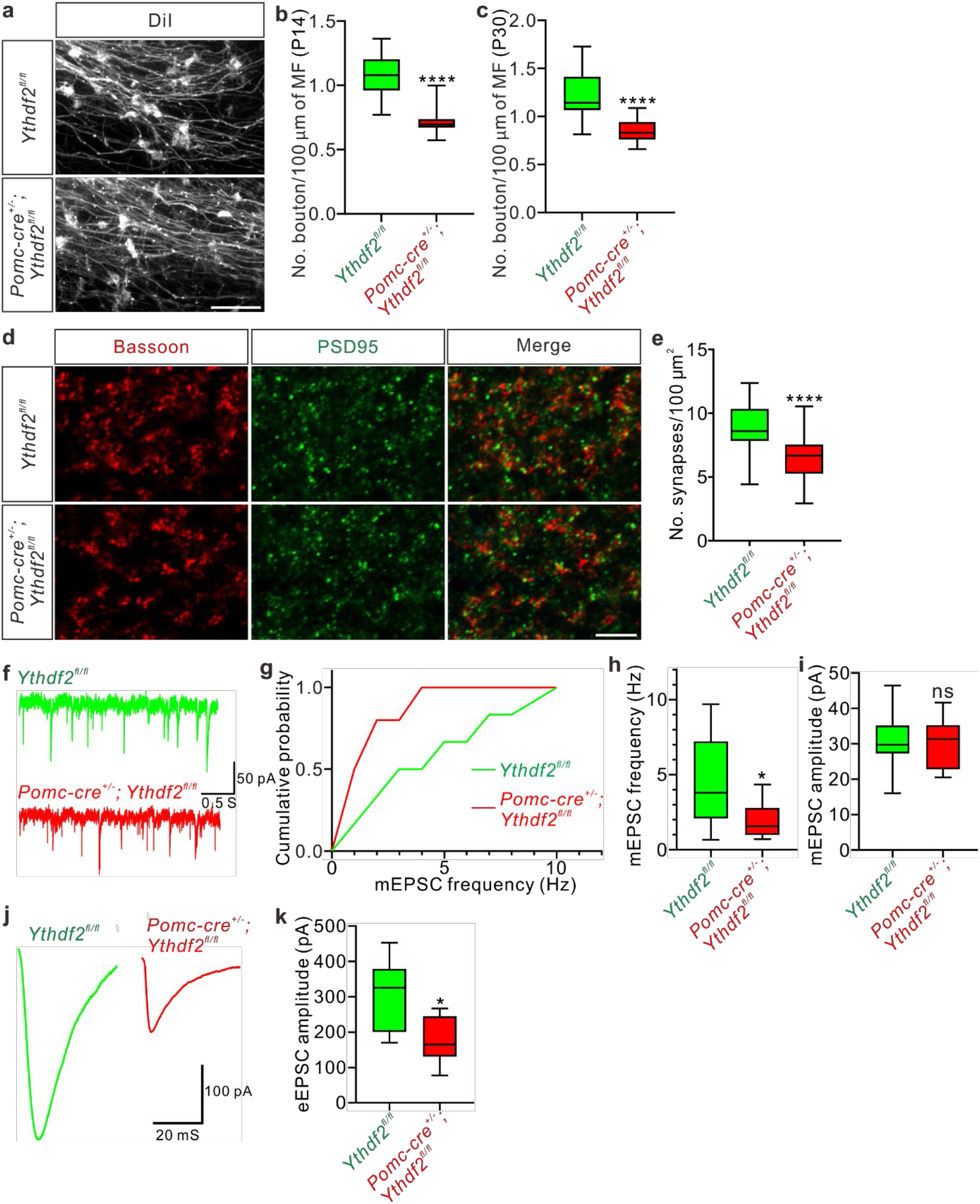
DG-CA3 synapse development and transmission are impaired in the DG-specific *Ythdf2* cKO mice. **a-c**, Bouton density on mossy fibers (MF) was decreased in the DG-specific *Ythdf2* cKO mice. DiI labeling was used to visualize mossy fibers and boutons in the stratum lucidum of *Ythdf2*^*fl/fl*^ and *Pomc-cre*^*+/-*^; *Ythdf2*^*fl/fl*^ mice at P14 (a). The numbers of MF boutons in *Pomc-cre*^*+/-*^; *Ythdf2*^*fl/fl*^ mice were reduced compared with those in *Ythdf2*^*fl/fl*^ mice at P14 (b) and P30 (c). Quantification of bouton numbers is represented as box and whisker plots: in b, *n* = 17 confocal fields for *Ythdf2*^*fl/fl*^, *n* = 18 confocal fields for *Pomc-cre*^*+/-*^; *Ythdf2*^*fl/fl*^, \*\*\*\**P* = 1.54E-6; in c, *n* = 20 confocal fields for *Ythdf2*^*fl/fl*^, *n* = 22 confocal fields for *Pomc-cre*^*+/-*^; *Ythdf2*^*fl/fl*^, \*\*\*\**P* = 2.84E-7; all by unpaired Student’s t-test. **d, e**, The MF-CA3 synapse numbers declined strikingly in the DG-specific *Ythdf2* cKO mice compared with control mice. Representative confocal images showed the excitatory synapses marked by co-staining of Bassoon and PSD95 in the stratum lucidum of the hippocampus at P60 (d). Quantification of synapse numbers is represented as box and whisker plots (e): *n* = 23 confocal fields for *Ythdf2*^*fl/fl*^, *n* = 28 confocal fields for *Pomc-cre*^*+/-*^; *Ythdf2*^*fl/fl*^, \*\*\*\**P* = 1.89E-6, by unpaired Student’s t-test. **f-i**, Representative traces (f), cumulative frequency distribution (g), and quantification of frequency (h) and amplitude (i) of mEPSCs in hippocampal CA3 pyramidal neurons of *Ythdf2*^*fl/fl*^ and *Pomc-cre*^*+/-*^; *Ythdf2*^*fl/fl*^ mice. The quantification is shown as box and whisker plots: *n* = 12 neurons for *Ythdf2*^*fl/fl*^, *n* = 10 neurons for *Pomc-cre*^*+/-*^; *Ythdf2*^*fl/fl*^, \**P* = 0.022 (h), *P* = 0.82 (i), ns, not significant, by unpaired Student’s t-test. **j, k**, Representative averaged traces (j) and quantification of amplitude (k) of eEPSCs evoked by stimulating the dentate gyrus granule cells. The quantification is shown as box and whisker plots: *n* = 6 synapses for *Ythdf2*^*fl/fl*^, *n* = 7 synapses for *Pomc-cre*^*+/-*^; *Ythdf2*^*fl/fl*^, \**P* = 0.020 (k), by unpaired Student’s t-test. At least 3 mice were analyzed for each genotype in each experiment. Scale bars, 20 μm (a), 5 μm (d).

These findings support that DG-specific *Ythdf2* knockout causes overgrowth of mossy fiber, reduction of MF boutons, and impairment of DG-CA3 synapse development and transmission.

### YTHDF2 destabilizes its target mRNAs to control DG granule cell axon growth

In order to elucidate the molecular mechanism by which YTHDF2 regulates DG mossy fiber growth and hippocampus-dependent learning and memory, we first carried out anti-YTHDF2 RNA immunoprecipitation (RIP) and sequencing (RIP-seq) to identify the target transcripts interacting with YTHDF2 in the hippocampus. 408 target mRNAs were identified (Supplementary Table 1). GO analysis demonstrated they were highly enriched in axon-related biological processes such as cell projection, axon, cell junction, and neuron projection (Fig. 5a), supporting the function of YTHDF2 to regulate axon growth at the molecular level.

**Fig. 5.**
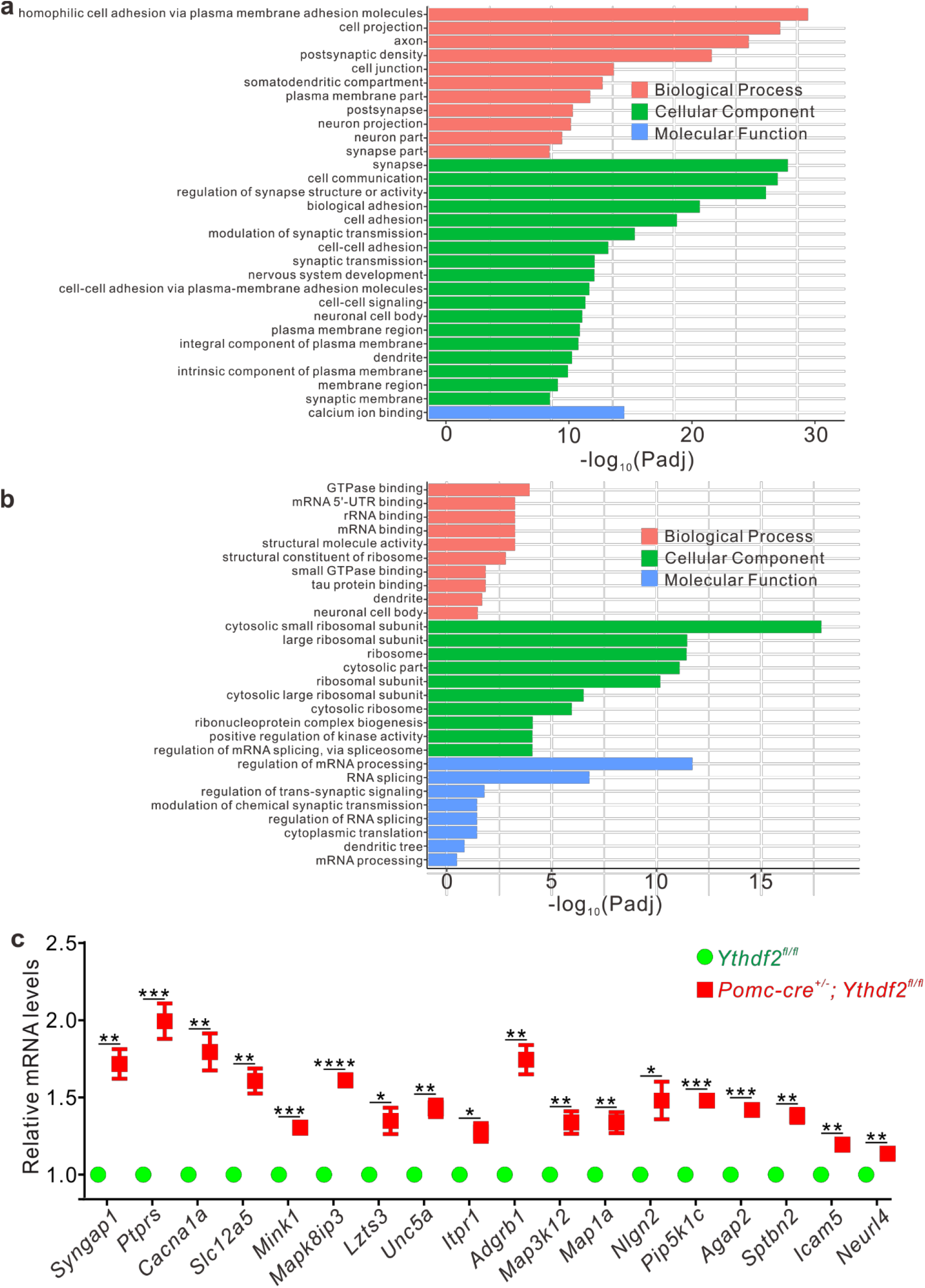
Target mRNAs of YTHDF2 were identified by integrating anti-YTHDF2 RIP-seq and RNAseq of DG-specific *Ythdf2* cKO. **a**, GO analysis of YTHDF2 target transcripts identified by anti-YTHDF2 RIPseq in P40 mouse hippocampus. **b**, GO analysis of transcripts with altered expression levels in DG-specific *Ythdf2* cKO dentate gyrus. **c**, Verification of upregulated target transcripts in DG-specific *Ythdf2* cKO mice by RT-qPCR. Data in c are represented as dot plot with mean ± SEM: \*\**P* = 0.0017 for *Syngap1*, \*\*\**P* = 9.47E-4 for *Ptprs*, \*\**P* = 0.0027 for *Cacna1a*, \*\**P* = 0.0018 for *Slc12a5*, \*\*\**P* = 3.73E-4 for *Mink1*, \*\*\*\**P* = 9.41E-6 for *Mapk8ip3*, \**P* = 0.015 for *Lzts3*, \*\**P* = 0.0015 for *Unc5a*, \**P* = 0.010 for *Itpr1*, \*\**P* = 0.0014 for *Adgrb1*, \*\**P* = 0.0094 for *Map3k12*, \*\**P* = 0.0076 for *Map1a*, \**P* = 0.017 for *Nlgn2*, \*\*\**P* = 2.01E-4 for *Pip5k1c*, ***P = 2.08E-4 for *Agap2*, \*\**P* = 0.0010 for *Sptbn2*, \*\**P* = 0.0034 for *Icam5*, \*\**P* = 0.0021 for *Neurl4*; *n* = 3 replicates; by unpaired Student’s t-test.

We next performed transcriptome profiling analysis of the DG-specific *Ythdf2* cKO mice. The DG subregion tissues were dissected from DG-specific *Ythdf2* cKO and their littermate control pups, and RNA sequencing was carried out to identify the transcripts that were changed in *Ythdf2*-ablated DG (cKO-RNAseq). Total 779 transcripts were identified with 536 upregulated and 243 downregulated (Supplementary Table 2). GO analysis showed that they were mostly enriched in RNA binding and processing (Fig. 5b).

By comparing the transcripts from RIP-seq and cKO-RNAseq, we identified 43 transcripts that bind YTHDF2 and display altered levels in DG ablated with *Ythdf2* (Supplementary Table 3). Among these transcripts, 37 were upregulated and 6 were downregulated in DG-specific *Ythdf2* cKO. The upregulation of 18 transcripts which are related to neuronal development or functions was further validated with RT-qPCR (Fig. 5c). We further focused on *Unc5a, Mapk8ip3*, and *Neurl4*. Unc5a was previously reported to induce neurite outgrowth ^30^, and Mapk8ip3 (also known as Jip3) was found to promote axon elongation in the hippocampal and cortical neurons ^31,32^, while there was no report on the function of Neurl4 in axon growth.

We examined whether Unc5a, Mapk8ip3, and Neurl4 mediated YTHDF2-regulated mossy fiber growth. We generated siRNAs against these transcripts, which led to efficient knockdown of corresponding mRNAs (Fig. 6a-c), and further caused decreases of GC axon growth (Fig. 6d-f). The m^6^A modification of *Unc5a, Neurl4*, and *Mapk8ip3* mRNAs was verified by anti-m^6^A pulldown (Fig. 6g). RNA half-life assay showed that the stability of these mRNAs was negatively controlled by YTHDF2 (Fig. 6h-j). YTHDF2 KD caused upregulation of Unc5a and Neurl4 protein levels (Supplementary Fig. 10). Importantly, knockdown of *Unc5a, Neurl4* or *Mapk8ip3* could efficiently rescue the GC axon overgrowth caused by DG-specific *Ythdf2* cKO (Fig. 6k-m).

**Fig. 6.**
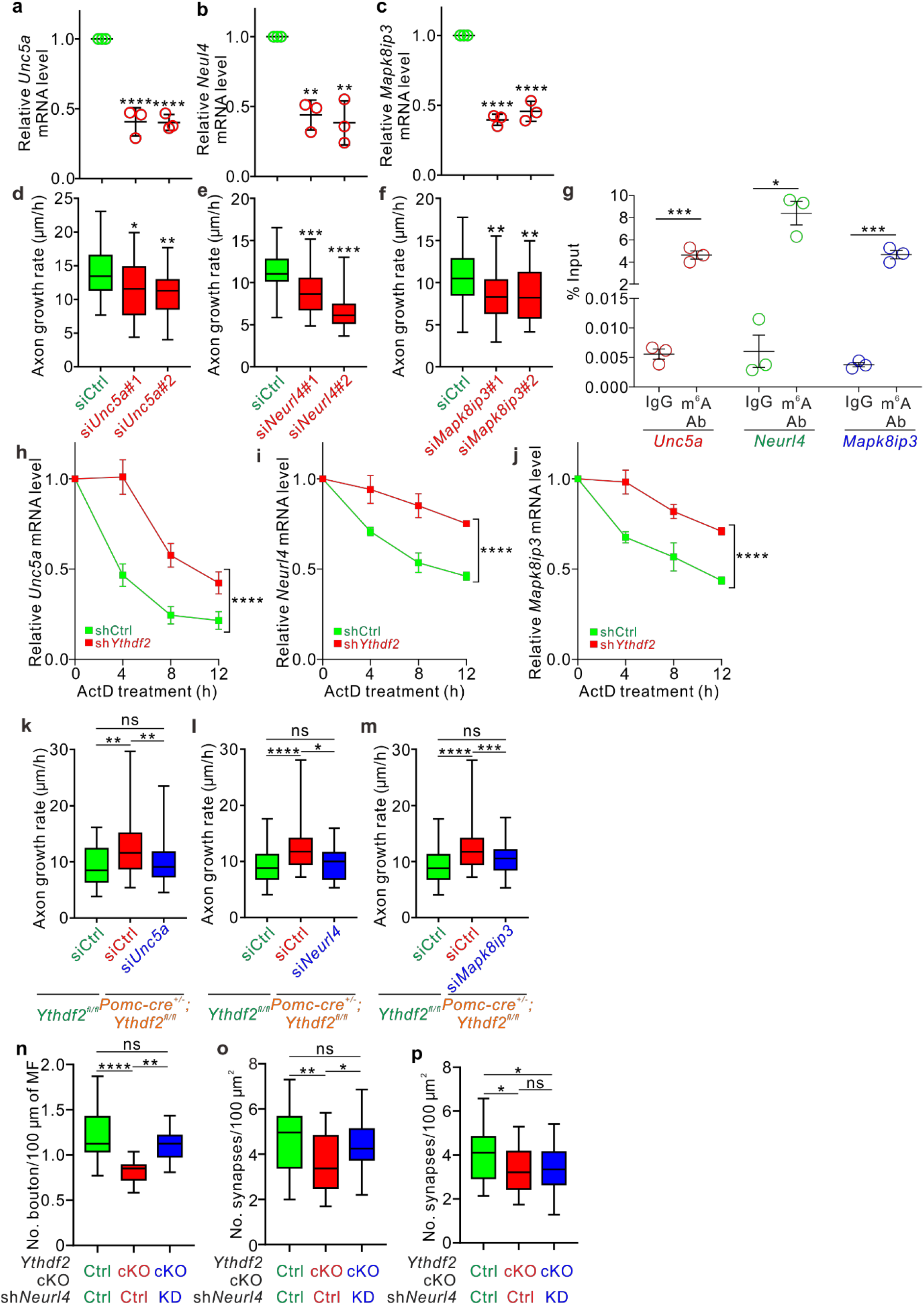
YTHDF2 destabilizes its m^6^A-modified target mRNAs to control DG granule cell axon growth. **a-c**, Knockdown efficiency of siRNAs against YTHDF2 target mRNAs were validated. GC neurons dissected from P6-8 DG were cultured and transfected with siRNAs against *Unc5a, Neurl4*, and *Mapk8ip3*, respectively. RNA was then extracted and RT-qPCR was carried out. **d-f**, Knockdown of *Unc5a, Neurl4*, or *Mapk8ip3* with siRNAs inhibited axon elongation of granule cells. **g**, m^6^A modification of *Unc5a, Neurl4, and Mapk8ip3* mRNAs was verified by anti-m^6^A pulldown followed by RT-qPCR. **h-j**, Half-lives of *Unc5a, Neurl4, and Mapk8ip3* mRNAs were increased after YTHDF2 knockdown. ActD, actinomycin D. **k-m**, Knockdown of *Unc5a, Neurl4, and Mapk8ip3* by siRNA attenuated the overgrowth of GC axons from DG-specific *Ythdf2* cKO mice. **n, o**, Knockdown of *Neurl4* by intracerebroventricular injection of shRNA rescued the decreases of bouton density and synapse numbers in DG-specific *Ythdf2* cKO mice. AAV-based sh*Neurl4* with GFP reporter was injected to DG at P0, and GFP-marked mossy fibers were analyzed at P14 to visualize MF boutons while PSD95/Bassoon IF was performed at P60 to analyze the MF-CA3 excitatory synapses. **p**, Knockdown of *Neurl4* by stereotaxic injections of shRNA at P30 could not rescue the decrease of synapse numbers in DG-specific *Ythdf2* cKO mice. Lentiviral sh*Neurl4* was injected to DG at P30, and PSD95/Bassoon IF was performed at P60 to analyze the MF-CA3 excitatory synapses. Data in a-c, g, and h-j are represented as dot plot with mean ± SEM, and all others are represented as box and whisker plots. (a-c) In a, *n* = 3 replicates, siCtrl vs si*Unc5a*#1, \*\*\*\**P* = 9.12E-6; siCtrl vs si*Unc5a*#2, \*\*\*\**P* = 8.67E-6. In b, *n* = 3 replicates, siCtrl vs si*Neurl4*#1, \*\**P* = 0.0019; siCtrl vs si*Neurl4*#2, \*\**P* = 0.0011. In c, *n* = 3 replicates, siCtrl vs si*Mapk8ip3*#1, \*\*\*\**P* = 1.07E-6; siCtrl vs si*Mapk8ip3*#2, \*\*\*\**P* = 2.07E-6. All by one-way ANOVA followed by Tukey’s post hoc test. (d-f) In d, siCtrl (*n* = 32 axons) vs si*Unc5a*#1 (*n* = 34 axons), \**P* = 0.010; siCtrl vs si*Unc5a*#2 (*n* = 34 axons), \*\**P* = 0.0025. In e, siCtrl (*n* = 41 axons) vs si*Neurl4*#1 (*n* = 38 axons), \*\*\**P* = 1.20E-4; siCtrl vs si*Neurl4*#2 (*n* = 41 axons), \*\*\*\**P* = 5.10E-9. In f, siCtrl (*n* = 45 axons) vs si*Mapk8ip3*#1 (*n* = 41 axons), \*\**P* = 0.0017; siCtrl vs si*Mapk8ip3*#2 (*n* = 45 axons), \*\**P* = 0.0018. All by one-way ANOVA followed by Tukey’s post hoc test. (g) In g, \*\*\**P* = 2.56E-5 for *Unc5a*, \**P* = 0.016 for *Neurl4*, \*\*\**P* = 2.80E-4 for *Mapk8ip3, n* = 3 replicates, by unpaired Student’s t-test. (h-j) \*\*\*\**P* = 1.02E-6 for h, \*\*\*\**P* = 3.05E-6 for i, \*\*\*\**P* = 2.53E-6 for j, *n* = 3 replicates, by two-way ANOVA followed by Tukey’s post hoc test. (k-m) In k, siCtrl with *Ythdf2*^*fl/fl*^ (*n* = 35 axons) vs siCtrl with *Pomc-cre*^*+/-*^; *Ythdf2*^*fl/fl*^ (*n* = 35 axons), \*\**P* = 0.0016; siCtrl with *Pomc-cre*^*+/-*^; *Ythdf2*^*fl/fl*^ vs si*Unc5a* with *Pomc-cre*^*+/-*^; *Ythdf2*^*fl/fl*^ (*n* = 32 axons), \*\**P* = 0.0099; siCtrl with *Ythdf2*^*fl/fl*^ vs si*Unc5a* with *Pomc-cre*^*+/-*^; *Ythdf2*^*fl/fl*^, *P* = 0.82. In l, siCtrl with *Ythdf2*^*fl/fl*^ (*n* = 44 axons) vs siCtrl with *Pomc-cre*^*+/-*^; *Ythdf2*^*fl/fl*^ (*n* = 48 axons), \*\*\*\**P* = 4.71E-6; siCtrl with *Pomc-cre*^*+/-*^; *Ythdf2*^*fl/fl*^ vs si*Neurl4* with *Pomc-cre*^*+/-*^; *Ythdf2*^*fl/fl*^ (*n* = 48 axons), \**P* = 0.013; siCtrl with *Ythdf2*^*fl/fl*^ vs si*Neurl4* with *Pomc-cre*^*+/-*^; *Ythdf2*^*fl/fl*^, *P* = 0.23. In m, siCtrl with *Ythdf2*^*fl/fl*^ (*n* = 44 axons) vs siCtrl with *Pomc-cre*^*+/-*^; *Ythdf2*^*fl/fl*^ (*n* = 48 axons), \*\*\*\**P* = 5.01E-6; siCtrl with *Pomc-cre*^*+/-*^; *Ythdf2*^*fl/fl*^ vs si*Mapk8ip3* with *Pomc-cre*^*+/-*^; *Ythdf2*^*fl/fl*^ (*n* = 47 axons), \*\*\**P* = 3.29E-4; siCtrl with *Ythdf2*^*fl/fl*^ vs si*Mapk8ip3* with *Pomc-cre*^*+/-*^; *Ythdf2*^*fl/fl*^, *P* = 0.84. ns, no significant; by one-way ANOVA followed by Tukey’s post hoc test. (n-p) In n, *Ythdf2*-cKO/shCtrl (*n* = 9 confocal fields) vs Ctrl/shCtrl (*n* = 24 confocal fields), \*\*\*\**P* = 5.16E-5; *Ythdf2*-cKO/sh*Neurl4* (*n* = 23 confocal fields) vs *Ythdf2*-cKO/shCtrl, \*\**P* = 0.0029. In o, *Ythdf2*-cKO/shCtrl (*n* = 29 confocal fields) vs Ctrl/shCtrl (*n* = 35 confocal fields), \*\**P* = 0.0025; *Ythdf2*-cKO/sh*Neurl4* (*n* = 31 confocal fields) vs *Ythdf2*-cKO/shCtrl, \**P* = 0.04. In p, *Ythdf2*-cKO/shCtrl (*n* = 20 confocal fields) vs Ctrl/shCtrl (*n* = 18 confocal fields), \**P* = 0.026; *Ythdf2*-cKO/sh*Neurl4* (*n* = 21 confocal fields) vs Ctrl/shCtrl, \**P* = 0.023. ns, no significant; by one-way ANOVA followed by Tukey’s post hoc test.

We further carried out in vivo rescue experiments by injections of AAV shRNA to the mouse brain. We focused on the novel player *Neurl4* which is not known to regulate axon development so far. AAV-based sh*Neurl4* with GFP reporter was intracerebroventricularly injected to DG at P0, and GFP-marked mossy fibers were analyzed at P14 to visualize MF boutons and PSD95/Bassoon IF was performed at P60 to analyze the MF-CA3 excitatory synapses. As shown in Fig. 6n, o, sh*Neurl4* could efficiently rescue the decrease of MF bouton density and synapse numbers in DG-specific *Ythdf2* cKO mice. To further verify that the phenotypes in bouton and synapse are consequences of GC axon overgrowth, we performed stereotaxic injection of AAV sh*Neurl4* to DG at P30 when GC axon growth is finished (thus, GC axon overgrowth in *Ythdf2* cKO could not be rescued), and analyzed the MF-CA3 excitatory synapses at P60. As shown in Fig. 6p, sh*Neurl4* injected at P30 could not rescue the decrease of synapse numbers in DG-specific *Ythdf2* cKO mice.

In summary, we identified a group of YTHDF2 target mRNAs which encode previously known and unknown axon growth-promoting proteins. Normally YTHDF2 destabilizes these m^6^A-modified target transcripts to control GC axon growth. In DG-specific *Ythdf2* cKO mice, *Neurl4, Unc5a*, and *Mapk8ip3* mRNAs were stabilized and upregulated accordingly, causing mossy fiber overgrowth in vivo, which further led to reductions in MF bouton density and synapse numbers.

## Discussion

Null knockout of the m^6^A reader YTHDF1 was shown to impair learning and memory in mice ^12^. In contrast to that study, our analysis argues against a vital role for YTHDF1 in hippocampus-dependent learning and memory as specific knockout of *Ythdf1* from dentate gyrus (DG), CA1 or CA3 does not show any defect in learning and memory. The possible explanation is that the global knockout of *Ythdf1* causes defects of universal circuits.

Our results demonstrate that only YTHDF2 in DG is required for the m^6^A-mediated learning and memory in the hippocampus. DG granule cells extend mossy fibers (MF) to form excitatory synapse with CA3 pyramidal neurons in the stratum lucidum ^33^. Deficits in MF-CA3 synapse can lead to intellectual disability ^34^. Alteration of MF-CA3 synapse was also found at an early stage in a mouse model of Alzheimer’s disease ^35^. Studies have shown that defects in DG or drugs working on DG could modulate MF-CA3 synaptic transmission and eventually impair hippocampal functions ^36^. Consistent with these, we find that DG-specific knockout of *Ythdf2* causes MF overgrowth, impairs MF-CA3 synapse formation and synaptic functions, and eventually causes defects in learning and memory.

We further find that YTHDF2 in DG regulates mossy fiber growth by controlling the stability of its m^6^A-modified target transcripts which encode proteins promoting axon growth and elongation. Among the three YTHDF2 targets identified in this study, Mapk8ip3 (Jip3) has been found to be concentrated in the distal axons of cultured hippocampus neurons ^31^. The axon guidance receptor Unc5a is also located in the growth cone of axons. Given that m^6^A modification has been shown to regulate local translation of mRNA in axons ^37^, it would be interesting to test if YTHDF2 works locally in axons and growth cones to control the expression of its targets to regulate axon growth and elongation. Neurl4 was not known to regulate axon development, and it would be interesting to further elucidate its working mechanisms.

In this study, we individually knocked out *Ythdf1* and *Ythdf2* from each hippocampal subregions respectively to find out that YTHDF2 in DG is the only m^6^A reader that mediates m^6^A modification in hippocampus-dependent learning and memory. Recent studies suggested that YTHDFs have overlapping and possibly redundant functions in mediating mRNA degradation by binding the same mRNAs and m^6^A sites ^38-40^, it would be interesting to see if the defects in hippocampus-dependent learning and memory are more pronounced if both *Ythdf2* and *Ythdf1* were depleted together.

## Materials and methods

### Animals

*Ythdf1*^*fl/fl*^ and *Ythdf2*^*fl/fl*^ mice were reported previously ^41,42^. *Pomc-cre* (Jax #010714), *Grik4-cre* (Jax #006474), *Camk2α-cre* (Jax #005359), *Rosa26-eYFP* (Jax #006148), and *Rosa26mT/mG* (Jax #007676) mice were all originally from the Jackson Laboratory stocks. All procedures or experiments using mice were performed following animal protocols approved by the Laboratory Animal Welfare and Ethics Committee of the Southern University of Science and Technology.

### Immunostaining

The immunostaining of primary neurons cultured on the cover glasses was done in accordance with the previous work ^41^.

The mouse brain sections for immunostaining were prepared from either freshly frozen brain tissues or perfused mice according to the needs in different experiments:

For freshly frozen tissue, mice were anesthetized with CO_2_ and decapitated, and the brains were rapidly embedded in O.C.T. compound (Sakura) and frozen immediately with dry ice. For immunostaining with m^6^A, YTHDF1, YTHDF2, Prox1, Elavl2 or WFS1 antibodies, freshly frozen and cut sections were air-dried, fixed with 4% PFA for 30 min at room temperature (RT), rinsed in PBS for 3 times with 5 min interval, and then blocked and permeabilized with PBS/10% normal donkey serum (NDS, Solarbio, SL050)/5% BSA/0.25% Triton x-100 at RT for 2 hr. For immunostaining with synaptic markers, the procedures followed a protocol described previously ^43^. Briefly, the freshly frozen tissue sections were fixed in cold methanol for 30 sec, rinsed with PBS twice with 5 min interval, and permeabilized in PBS/0.2% Triton x-100 for 20 min. After another twice-rinse by PBS, the slides were blocked with PBS/10% NDS at RT for 2 hr.

For perfusion, mice were perfused and the brains were taken out. Then the brains were fixed with 4% paraformaldehyde (PFA, Sigma) in 0.1 M phosphate buffer (PB) for overnight, dehydrated by 15% and 30% sucrose in 0.1 M PB overnight sequentially, and embedded in O.C.T. compound. 12-μm-thick cryosections were cut by a Leica CM1950 cryostat. The immunostaining with Calbindin, Calretinin, GFP, GFAP, Iba1, MBP, and SPO antibodies on perfused tissue sections generally followed the procedure of YTHDF1 staining, except that 4% PFA fixation was omitted here as the tissues had been fixed before. The immunostaining with BrdU antibody on perfused tissue sections was similar to Calbindin staining, except that prior to the staining steps, sections were first treated with 0.1 M sodium citrate at 95 °C for 30 min, followed by 2 M HCl treatment at RT for 30 min. After all these treatment, fixation or permeabilization steps, the slides were then incubated with primary antibodies at 4 °C for overnight, followed by incubation with secondary antibodies at RT for 1 hr. The slides were then mounted with mounting medium (Vector Laboratory, H-1200-10).

The sources and dilutions of antibodies are as following: m^6^A (Synaptic Systems, 202003, 1 μg/ml), YTHDF1 (Proteintech, 17479-1-AP, 1:1000), YTHDF2 (Proteintech, 24744-1-AP, 1:1000), GFP (Abcam, ab13970, 1:500), Prox1 (R&D Systems, AF2727, 0.2ng/ml), Elavl2 (Abclonal, A5918, 1:1000), WFS1 (Proteintech, 11558-1-AP, 1:1000), Bassoon (Enzo Life Sciences, ADI-VAM-PS003, 1:1000), PSD95 (Abcam, ab18258, 1 ng/ml), Calbindin (SWANT, 300, 1:200), Calretinin (Synaptic Systems, 214102, 1:200), BrdU (Abcam, ab6326, 1:200), Ki67 (Cell Signaling Technology, 12202S, 1:500), Neurl4 (Novus, NBP1-93574, 1:500), Unc5a (Proteintech, 22068-1-AP, 1:500), GFAP (Millipore, AB5541, 1:500), Iba1 (Millipore, MAPN92, 1:80), MBP (Invitrogen, MA5-15922, 1:100); Synaptoporin (SPO, SySy, 102 002, 1:500). Alexa Fluor-conjugated secondary antibodies (from Thermo or Jackson ImmunoResearch) were used (1:500 for 488 and 555, 1:200 for 647). Fluorescent images were captured using the laser-scanning confocal microscope Zeiss LSM800 with ZEN software. All images were captured with identical settings for different groups in the same experiment.

### m^6^A dot blot

The m^6^A dot blot was performed as described previously ^44^. In brief, RNA was extracted from brain tissues with FastPure^R^ Cell/Tissue Total RNA isolation Kit v2 (Vazyme, RC112-01) and loaded to the dried, positively charged Nylon membrane (Invitrogen). The m^6^A antibody (SySy, 202003, 1 μg/ml) and HRP-conjugated VHH anti-rabbit IgG (1:2500, KT Health, KTSM1322) were used in the study. After the exposure, the membrane was stained with 0.02% methylene blue (Shanghai Macklin Biochemical) for RNA loading control.

### BrdU injection

*Pomc-cre*^*+/-*^; *Ythdf2*^*fl/fl*^ mice and *Ythdf2*^*fl/fl*^ littermates received an injection of BrdU (Sigma) with the dose of 100 mg/kg intraperitoneally at P7. The brain tissues were collected 24 hr later.

### Behavioral studies

2-3-month-old male cKO and control mice were used in all behavioral experiments. Mice were habituated to the behavior room for 30 min before each day’s experiment. Behavioral tests were conducted at fixed time (13:00 to 17:00) by two operators blinding to the genotypes. The experimental area was cleaned with 75% ethanol before each test when the cleaning was applicable.

#### Morris water maze

Morris water maze assay was performed following the previously described steps ^45^ with some modifications. A circular pool with a depth of 50 cm and a diameter of 120 cm filled with water was divided into 4 quadrants, in which a platform was placed in one fixed quadrant. Visual cues were placed on the interior of the pool above the water surface. In the training experiments with a hidden platform, each mouse was placed into the water for 1 min, and the latency time was recorded as the mouse reached and stayed on the platform for 10 sec; if the mouse failed to find the platform in 60 sec, the operator would guide it to the platform and allow it to stay there for 10 sec before returning it to the cage, and record the latency as 60 sec. The average value from four trials, in which the same mouse was placed in water at different quadrants, was used to evaluate the performance of each mouse in the training experiments. In the probe test experiments, the platform was removed, and each mouse was given one probe trial for 60 sec. Two rounds of probe tests were conducted at day 3 and day 6, respectively. The movements of mice and the data were tracked, recorded, and analyzed by a cameral and the SMART 3.0 Small Animal Behavior Record Analysis System (RWD).

#### Open field

The mice were placed at the center of a square open area formed in a box (40 cm × 40 cm × 40 cm). Their activities within 10 min were recorded and analyzed using the SMART 3.0 Small Animal Behavior Record Analysis System (RWD).

#### Elevated plus-maze

The plus-maze placed 50 cm above the floor was composed of two closed arms, two open arms of the same size (50 cm × 9 cm) and a center area (9 cm × 9 cm). Mice were placed at the center area and allowed for 5 min to explore the place. Their activities were recorded and analyzed using the EthoVision software (Noldus).

#### Rotarod

Mice were placed on the rotarod apparatus (Ugo Basile, Stoelting) with a steadily accelerating rate of 4-40 rpm over 300 sec. The latency to fall of each mouse was recorded. The average value of three trials of each mouse was used for analysis.

#### Novel object recognition

Within the square area of a box (40 cm × 40 cm × 40 cm), two identical cylindrical volume blocks were placed diagonally on the floor. Mice were placed at the center of the square area and allowed for 10 min to explore and be familiar with the object. After an interval, one of the cylindrical volume blocks was replaced with a similar size square volume block at the same place, mice were then given another 10 min to explore the object. The activities of mice were recorded and analyzed using the EthoVision software (Noldus).

#### Fear conditioning

The fear conditioning test instrument (Anilab) comprises a transparent perspex chamber (21.5 cm × 17.1 cm × 16.2 cm) on a stainless-steel floor shock grid (floor bars are 3 mm in diameter and 6 mm in intervals) which is connected to a shocker-scrambler unit. During the experiment, the box was placed in a sound-poof box (60 cm × 50 cm × 60 cm). This test was completed in three days, where mice were given the tone cue-shock conditioning on the first day, their recall of contextual or tone-cued fear was tested on the second and the third day, consecutively. On the first day, mice were allowed to explore in the chamber for 2 min before receiving two pairings of a 30 sec tone cue (95 dB, 2.3 ± 0.3 kHz) co-terminating with a 2 sec foot shock (0.75 mA) at an intertrial interval (ITI) of 60 sec. 60 sec after the second tone-shock pairing, mice were sent back to their home cages. The freezing behavior of the mouse, which is defined as no less than one sec of no movement except respiration, was monitored and measured before and after the tone-shock pairing by AlsFC v1.0 (Anilab). On the second day, mice were placed in the chamber without tone cue or shock for 5 min, the freezing of mice during this period was monitored and used. On the third day, mice were placed in the chamber with a novel environment for 2 min followed by 2 min tone cue (95 dB, 2.3 ± 0.3 kHz), the freezing of mice during the pre-tone period and tones was monitored.

### Golgi staining

FD Rapid GolgiStain™ Kit (FD NeuroTechnologies; PK401) was used to examine the histology of dendrites and dendritic spines in *Ythdf2*^*fl/fl*^ and *Pomc-cre*^*+/-*^; *Ythdf2*^*fl/fl*^ littermates at P31 and P61. After mice were anesthetized with CO_2_ and decapitated, the brain was rapidly removed and subjected to the procedures instructed by the kit manual. Sections with a thickness of 100 μm were cut on a vibrating microtome (Leica VT1200S). The laser-scanning confocal microscope Nikon A1R with NIS software was used for imaging DG dendrites with multiple z-stacks. The Olympus BX53 microscope with a DP73 camera and cellSens Standard software was used for capturing and analyzing the images of DG dendritic spines. The quantification was done with ImageJ.

### Neuron culture

Unless otherwise specified, all reagents used for neuron culture were from Thermo Fisher Scientific. Dissection and culturing of DG granule cells from P6-8 mouse pups were generally following the procedures reported previously ^46^ with some changes. Briefly, DG was dissected in Hibernate-A medium, and then digested by HBSS/0.1% Trypsin (Sigma)/0.1% DNase I (Sigma). The dissociated GC neurons were then suspended with the plating medium containing Neurobasal-A, 10% FBS, GlutaMAX-1 (1×) and penicillin-streptomycin (1×). For the axon growth assay, the GC neurons were pipetted into the soma compartments of the microfluidic chambers coated with poly-D-lysine (R&D). After 0.5-1 hr, when the neurons were attached, the culturing medium composed of Neurobasal-A, B27 (1×), GlutaMAX1 (1×) and penicillin-streptomycin (1×) was added to the chambers. For other experiments without using the microfluidic chambers, GC neurons were plated to the 24-well plates coated with poly-D-lysine with plating medium, which was replaced with culturing medium 4 hr later.

### Knockdown and overexpression using the lentiviral system

pLKO.1-TRC (Addgene, 10878) vector was used for the knockdown of YTHDF2. The target sequences of shRNA are as following: shCtrl, 5’-GCATCAAGGTGAACTTCAAGA-3’; sh*Ythdf2*#1, 5’-GCTCCAGGCATGAATACTATA-3’; sh*Ythdf2*#3, 5’-GGACGTTCCCAATAGCCAACT-3’. YTHDF2 overexpression construct was made similarly as YTHDF1 which was reported previously ^41^. The generation and purification of lentivirus were done as previously reported ^37^. The samples were collected or analyzed 48 hr after the infection.

### Knockdown using the AAV system in vivo

Adeno-associated virus (AAV) of serum type 8 was used for knockdown of *Neurl4* in vivo. The target sequences of shRNA are as follows: shCtrl, 5’-TTCTCCGAACGTGTCACGTAA-3’; sh*Neurl4*, 5’-GCTTGCTCTTCCATCCTAA-3’. The plasmid construction and the virus package were done by Hanbio Biotechnology Co. Ltd. The AAV (1.2 × 10^12^ viral genome/ml) was either intracerebroventricularly injected into the neonatal mouse brain or delivered to the DG by stereotaxic intracranial injection according to the aims of different experiments with procedures adapted from the previously published works ^47,48^. In brief, for intracerebroventricular injection of AAV into the neonatal mouse brain, the mice were cryoanesthetized with aluminum foil surrounded by ice, and 2 μl virus was injected into each lateral ventricle using 10 μl Hamilton syringes with a depth of 1.5 mm. The mice were then returned to the home cage after they had recovered entirely on a warming blanket. The mice were then sacrificed at P14 for the examination of mossy fiber and boutons, or at P60 for examination of synapses. For the stereotaxic injection, P30 mice were anesthetized and fixed in stereotaxic apparatus with their fur on the skull shaved, and the skin was cleaned with 70% ethanol. Afterward, the skull over the target area was thinned, 0.5 μl virus was then injected with an injection micropipette. The injection position was cautiously calculated to make sure that the virus was delivered to the DG subregion. The mice were then sent back to the home cage after they recovered from anesthesia. The mice were sacrificed at P60 for examination of synapses.

### Knockdown using siRNA

siRNA-mediated knockdown of *Neurl4, Unc5a*, and *Mapk8ip3* was carried out with GeneSilencer siRNA Transfection Reagent (Genlantis) following the previously reported protocol ^37^. The target sequences of siRNA are as following: siCtrl, 5’-UUCUCCGAACGUGUCACGUTT-3’; si*Unc5a*#1, 5’-CCAGGUCGAUCACGUCAUUTT-3’; si*Unc5a*#2, 5’-GCUGACUCAUCCAUCCUUATT-3’; si*Neurl4*#1, 5’-GCUUGCUCUUCCAUCCUAATT-3’; si*Neurl4*#2, 5’-GCAAUACGAUGCGCAACAATT-3’; si*Mapk8ip3*#1, 5’-AGCGUCCCACCUCUCUGAAUGUCUU-3’; si*Mapk8ip3*#2, 5’-CCCAGGGAAUUGUAAACAATT-3’.

### Western blotting

Hippocampal tissues from different stages of wild type (WT) mouse brains, or cultured neurons infected with lentivirus carrying shCtrl, sh*Ythdf2*#1 or sh*Ythdf2*#3 for 48 hr, were collected using RIPA buffer with cOmplete Protease Inhibitor Cocktail (Roche). The YTHDF1 or YTHDF2 protein level was measured by Western blotting using a YTHDF1 (Proteintech, 17479-1-AP, 1:1000) or YTHDF2 antibody (Proteintech, 24744-1-AP, 1:1000).

### Axon growth assay

Dissociated GC neurons from WT mice or mice with different genotypes were plated to the soma compartments of the microfluidic chambers. Lentivirus harboring shRNA or siRNAs were infected or transfected according to the specific needs of each experiment. 48 hr later, the bright-field images of the axonal compartment were captured at two different time points with an inverted microscope Nikon LS100. Axons were manually traced and measured using ImageJ.

### DiI labeling of mossy fibers

P14 and P30 mice were perfused with 4% PFA/0.1 M PB, and the brains were taken out and fixed at 4 °C for overnight. 200-μm-thick sections were cut on a vibratome (Leica VT1200S) and stored in PBS. DiI crystals (Thermo Fisher Scientific, D282) were placed on the hilar region of the dentate gyrus, and the sections were then kept at RT for 24 hr before being mounted with the mounting medium (Vector Laboratory, H-1000-10). Images for DiI fluorescence were acquired with a LSM800 confocal microscopy (Zeiss) with ZEN 2.3 software. The length of DiI labeled MF axons and the size of MF boutons were measured using ImageJ. The number of boutons was counted manually in fields-of-interest and was divided by the total length of MF axons in that field.

### Electrophysiological recording of hippocampal slices

Whole-cell recordings were carried out on the transverse hippocampal slices (350-μm-thick) which were prepared from P21-P28 cKO and control mice according to the previously published literature ^49^. Briefly, mice were deeply anesthetized using isoflurane and decapitated. Slices were cut using a vibratome (Campden 7000smz-2) in the ice-cold cutting solution containing 64 mM NaCl, 25 mM NaHCO_3_, 2.5 mM KCl, 1.25 mM NaH_2_PO_4_, 10 mM glucose, 120 mM sucrose, 0.5 mM CaCl_2_, and 7 mM MgCl_2_. Slices were then incubated at 35 °C for ∼30 min, and subsequently transferred to room temperature in the storage solution containing 87 mM NaCl, 25 mM NaHCO_3_, 2.5 mM KCl, 1.25 mM NaH_2_PO_4_, 10 mM glucose, 75 mM sucrose, 0.5 mM CaCl_2_, and 7 mM MgCl_2_ (gassed with 95% O_2_ and 5% CO_2_). Fifteen minutes before recording, slices were transferred to the recording chamber and superfused with ACSF (artificial cerebrospinal fluid) containing 125 mM NaCl, 25 mM NaHCO_3_, 2.5 mM KCl, 1.25 mM NaH_2_PO_4_, 2 mM CaCl_2_, and 1 mM MgCl_2_, gassed with 95% O_2_ and 5% CO_2_. CA3 pyramidal neurons were whole-cell voltage-clamped at -70 mV using an EPC10/2 amplifier with glass pipettes (2-4 MΩ). The patch pipettes were filled with the intracellular solution containing 130 mM K-gluconate, 2 mM KCl, 2 mM MgCl_2_, 2 mM Na_2_ATP, 10 mM HEPES, 10 mM EGTA (pH 7.2 with KOH, osmolarity 310–315 mOsm). Series resistance was carefully monitored and typically below 15 MΩ, and only recordings with stable series resistance were included in the analysis. Miniature excitatory postsynaptic currents (mEPSCs) were recorded in the presence of 1 μM TTX. Evoked EPSCs (eEPSCs) were recorded by stimulating the dentate gyrus granule cells with a stimulating electrode (Microprobe CEA200).

### RNA immunoprecipitation and sequencing (RIP-seq)

Hippocampal tissues from P40 WT mice were collected and lysed with IP lysis buffer in EZ-Magna RIP™ RNA-Binding Protein Immunoprecipitation Kit (Merck-Millipore). YTHDF2 (Proteintech, 24744-1-AP) and control IgG were used to pull down the transcripts following the kit manual. The eluted RNA samples were then used for cDNA library construction, sequencing, and analysis. The enrichment of RNAs with YTHDF2 IP was normalized to IgG. Sequencing results in each group were from two independent replicates.

### Transcriptome profiling by RNA sequencing

Total RNAs were isolated from the dentate gyrus of P40 *Ytdhdf2*^*fl/fl*^ and *Pomc-cre*^*+/-*^; *Ythdf2*^*fl/fl*^ hippocampi using TRIzol Reagent (Thermo Fisher Scientific). The mRNA purification, cDNA library construction, sequencing and analyzing were done following standard procedures. The gene expression profiling results were from three independent replicates.

### Anti-m^6^A immunoprecipitation

Total RNA was extracted from the dentate gyrus of the hippocampus in P6∼P8 WT mice. Immunoprecipitation of m^6^A-modified transcripts was carried out with Magna MeRIP™ m^6^A kit (Merck-Millipore) following the manual. m^6^A antibody (Synaptic Systems, 202003) and corresponding control IgG were used in this experiment. The RNA samples pulled down from the experiment were then subject to RT-qPCR.

### RNA stability assay

GC neurons infected with lentiviral sh*Ythdf2* and control shRNA for 48 hr were treated with actinomycin D (ActD, 5 μg/ml, ApexBio Technology) and collected at indicated time points. The mRNA levels of *Unc5a, Neurl4*, or *Mapk8ip3* in different conditions were normalized to the 18s ribosome RNA, before comparing the different groups.

### RT-qPCR

Total RNA was extracted using TRIzol Reagent (Thermo Fisher Scientific). Reverse transcription with 1 μg of total RNA was conducted using PrimeScript^™^ RT Master Mix (Takara) following the manual. Synthesized cDNA was then subjected to real-time quantitative PCR using 2×ChamQ^™^ Universal SYBR qPCR Master Mix (Vazyme) with a BioRad qPCR CFX 96 machine. The primers used in m^6^A IP and RNA stability assay are as following: 5’-TCTTCAAGTGCAACGGGGAA-3’ and 5’-CATGGTTGGCAATCCGCTG-3’ for *Unc5a*; 5’-ATACGTGGATGCTGAGTGGTA-3’ and 5’-GCCGGTAGCATTGGTGATAG-3’ for *Neurl4*; 5’-CAGCGGCACACAGAGATGA-3’ and 5’-ACATTCAGAGAGGTGGGACG-3’ for *Mapk8ip3*; 5’-GCTTAATTTGACTCAACACGGGA-3’ and 5’-AGCTATCAATCTGTCAATCCTGTC-3’ for 18s rRNA ^50^. The primers used for verifying siRNA efficiency are as following: 5’-GTGCAACGGGGAATGGGT-3’ and 5’-AGTTCTTGCGCAAATAGGCAAT-3’ for si*Unc5a*#1; 5’-CTCTGCCTGCACACCTCTTC-3’ and 5’-GCAGATGGGGATTGTCTGCT-3’ for si*Unc5a*#2; 5’-ATGGCATCGATCAGGGCGTA-3’ and 5’-GCGTCCTTCGTGGGTGATAG-3’ for si*Neurl4*#1; 5’-ATACGTGGATGCTGAGTGGTA-3’ and 5’-GCCGGTAGCATTGGTGATAG-3’ for si*Neurl4*#2; 5’-AGTCGTGGAGGAAAAGCAGGA-3’ and 5’-TACGGCTATTGCGCTTCTCT-3’ for si*Mapk8ip3*#1; 5’-AACGTCAGTATCGGCATGGG-3’ and 5’-GCGCACTGAAAACTCCCCTA-3’ for si*Mapk8ip3*#2; 5’-CAAGGAGTAAGAAACCCTGGAC-3’ and 5’-GGATGGAAATTGTGAGGGAG-3’ for *Gapdh*. The primers used for measuring m^6^A writer/eraser/reader mRNA levels are as following: 5’-GAAGAGGAGGTGGTGCGTAAG-3’ and 5’-CGGAGACAGCACCAAGCATAC-3’ for *Ythdf1*, 5’-GAAGAGGAGGTGGTGCGTAAG-3’ and 5’-CGGAGACAGCACCAAGCATAC-3’ for *Ythdf2*, 5’-GAAGAGGAGGTGGTGCGTAAG-3’ and 5’-CGGAGACAGCACCAAGCATAC-3’ for *Ythdf3*, 5’-GAAGAGGAGGTGGTGCGTAAG-3’ and 5’-CGGAGACAGCACCAAGCATAC-3’ for *Mettl3*, 5’-GAAGAGGAGGTGGTGCGTAAG-3’ and 5’-CGGAGACAGCACCAAGCATAC-3’ for *Mettl14*, 5’-GAAGAGGAGGTGGTGCGTAAG-3’ and 5’-CGGAGACAGCACCAAGCATAC-3’ for *Alkbh5*, 5’-GAAGAGGAGGTGGTGCGTAAG-3’ and 5’-CGGAGACAGCACCAAGCATAC-3’ for *Fto*.

### Statistical analysis

All experiments were conducted with at least three independent biological replicates. For in vitro experiments and in vivo tissue analysis, no statistical methods were used to determine sample size. We collected the samples as much as we can, which were no less than those in the literatures in the field. The sample sizes in animal studies were chose according to related literatures in the field. The maximum and minimum values were excluded in the behavior analysis, which was a pre-established criterion. No exclusions were done in other analysis. For in vitro experiments using primary neurons from wild type (WT) mice, the WT mice of mentioned stages were randomly allocated in primary culture experiments. No randomization was used in phenotype observation experiments. The investigators were blind to the genotypes of mice in the behavior tests and the DiI tracing experiments of mossy fiber. Quantification data are represented either in box and whisker (25^th^ -75^th^ percentiles for boxes, minimum and maximum for whiskers, medians for horizontal lines) or in dot blots with mean ± SEM. Statistical analysis was performed with SPSS software, and the graphs were made using Graphpad Prism 9. Unpaired two-tailed Student’s t-test was used to compare the values of two groups. One-way analysis of variance (ANOVA) with Tukey’s Honest Significant Differences (Tukey’s HSD) post hoc test was used to compare changes in three groups. Two-way ANOVA with Tukey’s HSD post hoc test was used to compare values in two groups with different time points. The variances between groups were tested and were comparable.

## Data availability

RNAseq of *Ythdf2* cKO in DG and anti-YTHDF2 RIPseq data have been deposited in NCBI’s Gene Expression Omnibus (GEO) and are accessible through accession numbers GSE171790 and GSE171791.

## Supporting information

Supplemental fiigures

## Acknowledgements

We thank Xin Ren for technical assistance, and other members of Ji and Dong laboratories for help, technical support, and comments on the manuscript. We thank Yu Chung Tse and Yilin Wang of SUSTech Core Research Facilities for their help in imaging and analysis of DG dendrites and spines after Golgi staining. We thank Prof. Shengtao Hou for support in the behavioral tests. We thank Prof. Jun Xia at The Hong Kong University of Science and Technology (HKUST) for co-mentoring the Ph.D. student in the SUSTech-HKUST Joint Ph.D. Program. This work was supported by National Natural Science Foundation of China (31871038 and 32170955 to S.-J.J., 31871031 to W.D.), Shenzhen-Hong Kong Institute of Brain Science-Shenzhen Fundamental Research Institutions (2022SHIBS0002), High-Level University Construction Fund for Department of Biology (internal grant no. G02226301), Science and Technology Innovation Commission of Shenzhen Municipal Government (ZDSYS20200811144002008), Ministry of Science and Technology of the People’s Republic of China (2019YFE0120600), and Department of Science and Technology of Sichuan Province (2019YJ0481).

## Author Contributions

S.-J.J. conceived the idea and designed the experiments. M.Z. and P.H. performed and analyzed the experiments with help of P.C. X.G. and C.L. performed the electrophysiology experiments. F.L. and Z.Z. performed the behavioral tests. F.L., L.Y. and J.Y. performed the RIPseq and RNAseq analysis. M.Z., S.-J.J. and W.D. wrote the manuscript with input from all authors.

## Conflict of interest

The authors declare that they have no conflict of interest.

## Figures and figure legends

**Supplementary Table 1. List of YTHDF2 target mRNAs by anti-YTHDF2 RIPseq of the hippocampus**.

**Supplementary Table 2. Transcriptome analysis of dentate gyrus from DG-specific *Ythdf2* cKO hippocampus**

**Supplementary Table 3. Overlapping transcripts in anti-YTHDF2 RIPseq and DG-*Ythdf2*-cKO-RNAseq**

## Notes

### Competing Interest Statement

The authors have declared no competing interest.

