## Supplemental fiigures for "YTHDF2 in dentate gyrus is the m^6^A reader mediating m^6^A modification in hippocampus-dependent learning and memory"

**This file includes Supplementary figures 1 to 10.**

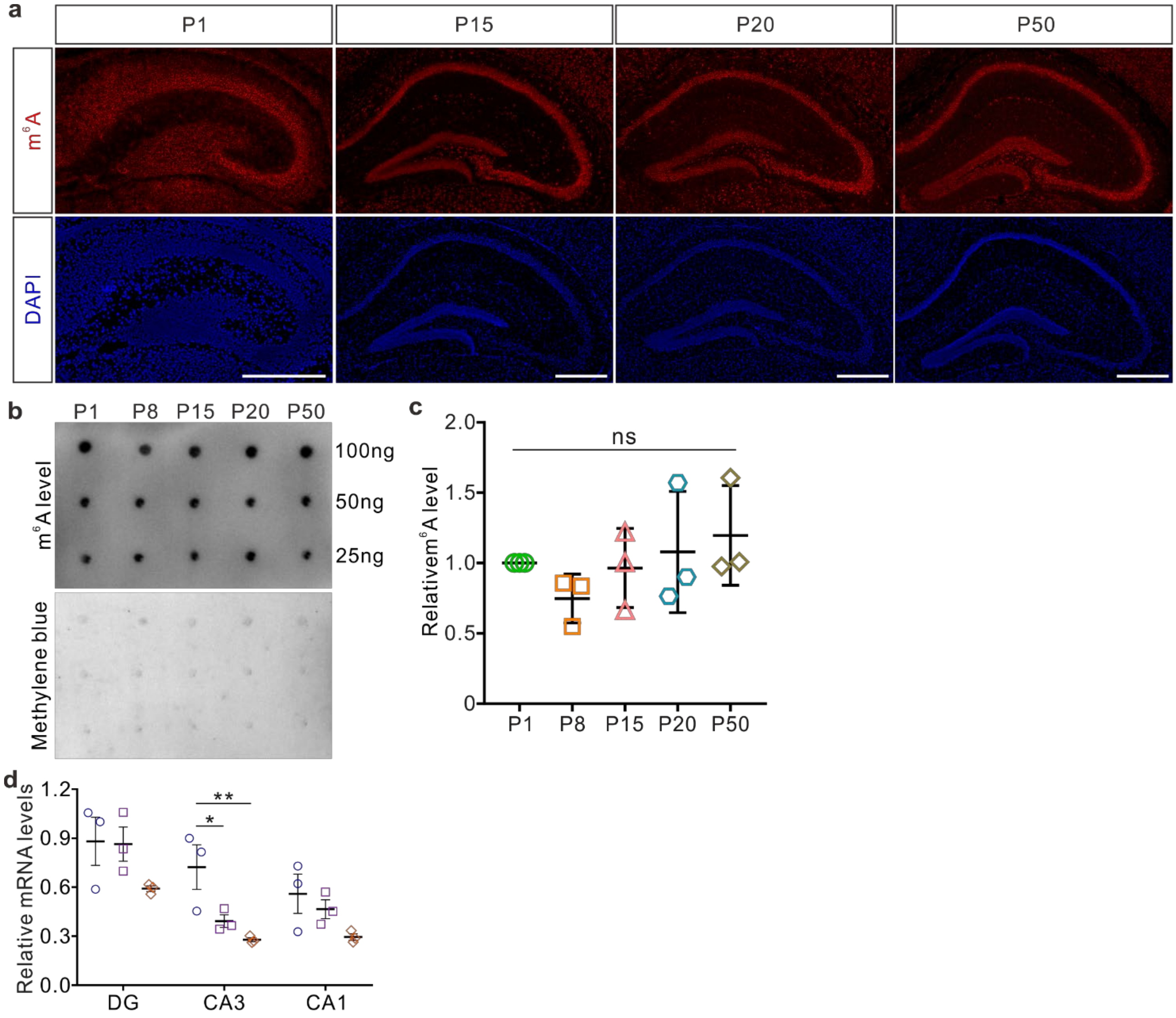


**Supplementary Fig. 1. High enrichment of m$^6$A in the hippocampus by anti-m$^6$A immunofluorescence.**

**a,** Representative confocal images of anti-m$^6$A immunofluorescence (IF) of mouse coronal brain sections showed strong m$^6$A signals in the hippocampi of P1, P15, P20, and P50 mice. **b, c,** Anti m$^6$A dot blots of P1-P50 WT hippocampi. The amount of RNAs used in the assay is indicated. Methylene blue staining was used as the loading control. Quantification of dot blots showed that the m$^6$A modification levels in the hippocampus are not changed in the shown early postnatal stages (c). **d,** Relative expression levels of *Ythdf1*, *Ythdf2*, and *Ythdf3* in different subregions of the hippocampus by RT-qPCR. All quantifications are represented in dot plots as mean ± SEM. In c, $n$ = 3 mice for each stage. In d, $n$ = 3 mice for each subregion at P8: for CA3 subregion, *Ythdf1* vs *Ythdf2*, *$P$ = 0.03; *Ythdf1* vs *Ythdf3*, **$P$ = 0.0087; unless indicated by * showing significantly different, all other comparisons with $P$ values “not significant” (ns) were not shown; all by one-way ANOVA followed by Tukey’s post hoc test. Scale bars, 500 μm.

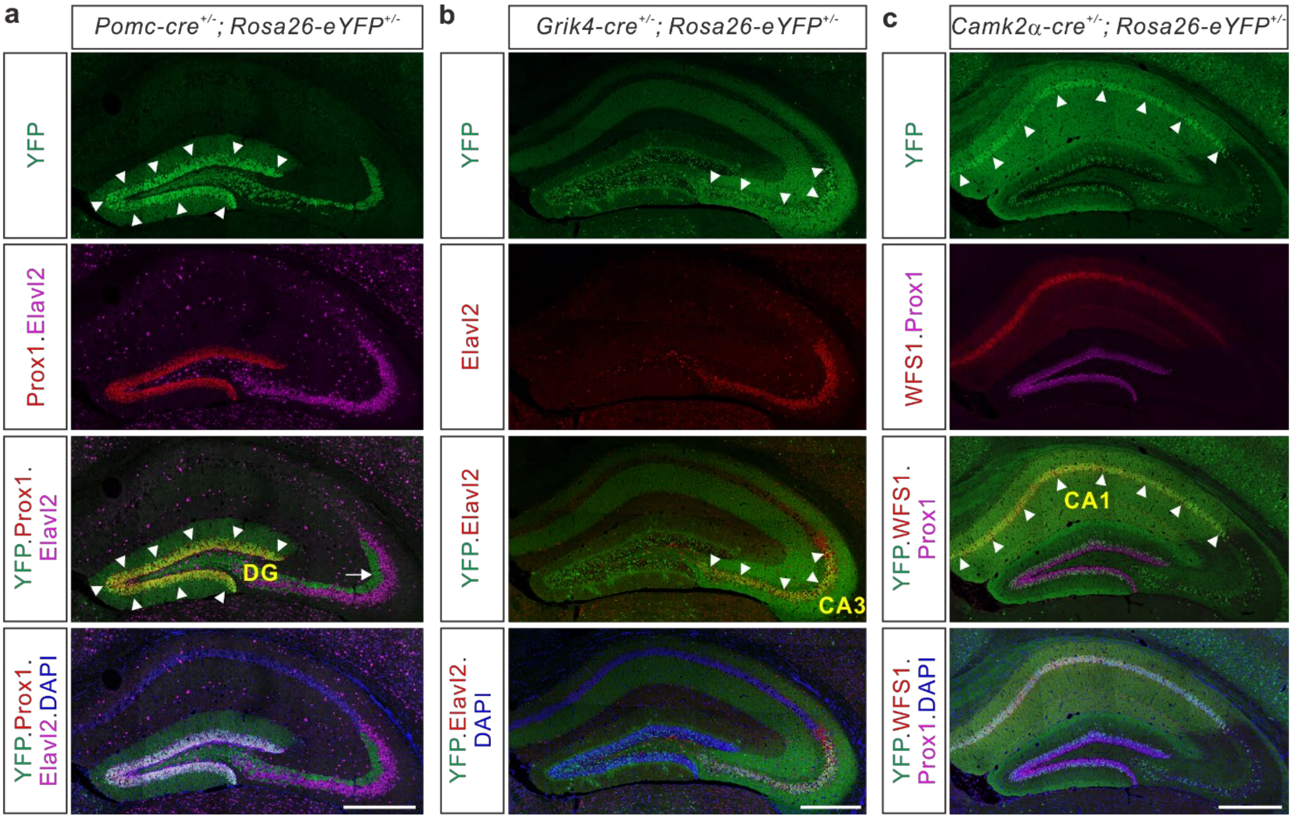


**Supplementary Fig. 2. DG-, CA3-, and CA1-specific Cre lines were validated by crossing with the *Rosa26-eYFP* reporter.**

**a,** In P20 *Pomc-cre*$^{+/-}$; *Rosa26-eYFP*$^{+/-}$ mice, Cre expression indicated by YFP IF was detected exclusively in the Prox1-labelled DG (white arrowheads) granule cells. Notice that YFP-labelled mossy fiber projected to the stratum lucidum (SL, white arrow) of the CA3 marked by Elavl2. **b,** Robust YFP signals indicated restricted Cre expression in CA3 (white arrowheads) pyramidal neurons marked by Elavl2 in P32 *Grik4-cr*$^{+/-}$*e; Rosa26-eYFP*$^{+/-}$ mice. Scattered YFP signals were detected in the DG area. **c,** CA1 (white arrowheads) pyramidal neurons marked by WFS1 in P26 *Camk2α-cre*$^{+/-}$*; Rosa26-eYFP*$^{+/-}$ mice were densely labelled with YFP. Scattered YFP signals were also found in DG and CA3 areas. Scale bars, 500 μm (a-c).

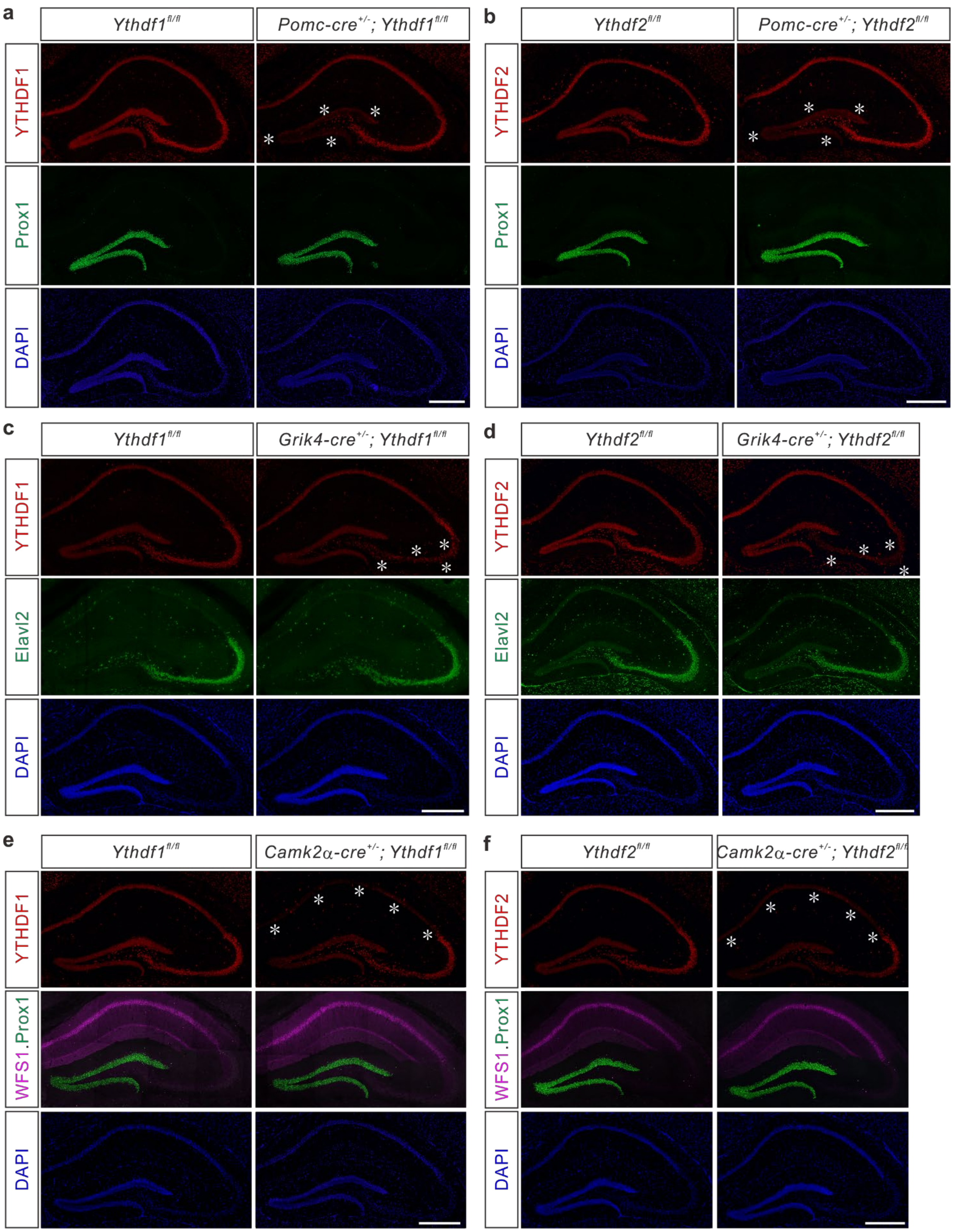


**Supplementary Fig. 3. *Ythdf1* and *Ythdf2* are specifically ablated from DG, CA3, and CA1, respectively, using hippocampal subregion-specific Cre lines.**

**a, b,** *Pomc-cre*$^{+/-}$*; Ythdf1*$^{fl/fl}$ and *Pomc-cre*$^{+/-}$*; Ythdf2*$^{fl/fl}$ are DG-specific *Ythdf1* and *Ythdf2* cKO, respectively. **c, d,** *Grik4-cre*$^{+/-}$*; Ythdf1*$^{fl/fl}$ and *Grik4-cre*$^{+/-}$*; Ythdf2*$^{fl/fl}$ are CA3-specific *Ythdf1* and *Ythdf2* cKO, respectively. **e, f,**

*Camk2α-cre*$^{+/-}$*; Ythdf1*$^{fl/fl}$ and *Camk2α-cre*$^{+/-}$*; Ythdf2*$^{fl/fl}$ are CA1-specific *Ythdf1* and *Ythdf2* cKO, respectively. YTHDF1 (a, c, e) and YTHDF2 (b, d, f) are specifically and efficiently knocked out from adult hippocampal DG (a, b), CA3 (c, d), and CA1 (e, f) regions, respectively. Prox1, Elavl2, and WFS1 are the markers for DG, CA3, and CA1, respectively. The asterisks highlight the efficient knockout in the corresponding hippocampal subregions. Scale bars, 500 μm.

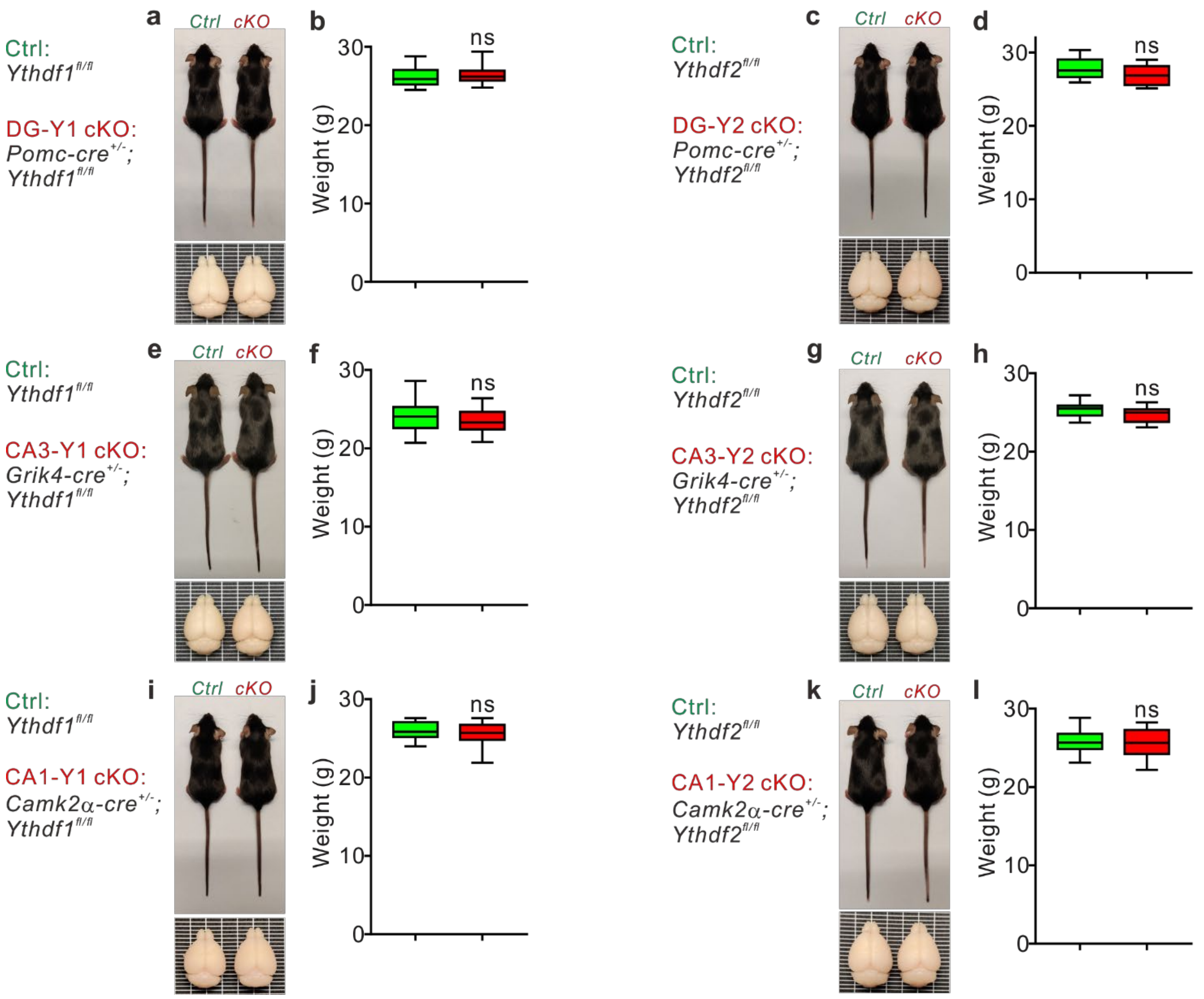


**Supplementary Fig. 4. Hippocampal subregion-specific *Ythdf1* or *Ythdf2* cKO mice have normal body size and brain morphology.**

DG-, CA3-, and CA1-specific *Ythdf1* or *Ythdf2* cKO mice, designated as DG-Y1 cKO (a, b), DG-Y2 cKO (c, d), CA3-Y1 cKO (e, f), CA3-Y2 cKO (g, h), CA1-Y1 cKO (i, j), and CA1-Y2 cKO (k, l), have normal body size and brain morphology (a, c, e, g, i, k), and body weight (b, d, f, h, j, l) compared with control mice (*Ythdf1$^{fl/fl}$* or *Ythdf2$^{fl/fl}$*, designated as *Ctrl*). Quantification of body weights is represented as box and whisker plots. For DG-Y1 cKO, $n$ = 15 mice for each genotype in b; for DG-Y2 cKO, $n$ = 14 mice for Ctrl and $n$ = 10 mice for DG-Y2 cKO in d; for CA3-Y1 cKO, $n$ = 18 mice for each genotype in f; for CA3-Y2 cKO, $n$ = 16 mice for Ctrl and $n$ = 14 mice for CA3-Y2 cKO in h; for CA1-Y1 cKO, $n$ = 10 mice for each genotype in j; for CA1-Y2 cKO, $n$ = 12 mice for Ctrl and $n$ = 13 mice for CA1-Y2 cKO in l; ns, not significant; all by unpaired Student's t-test.

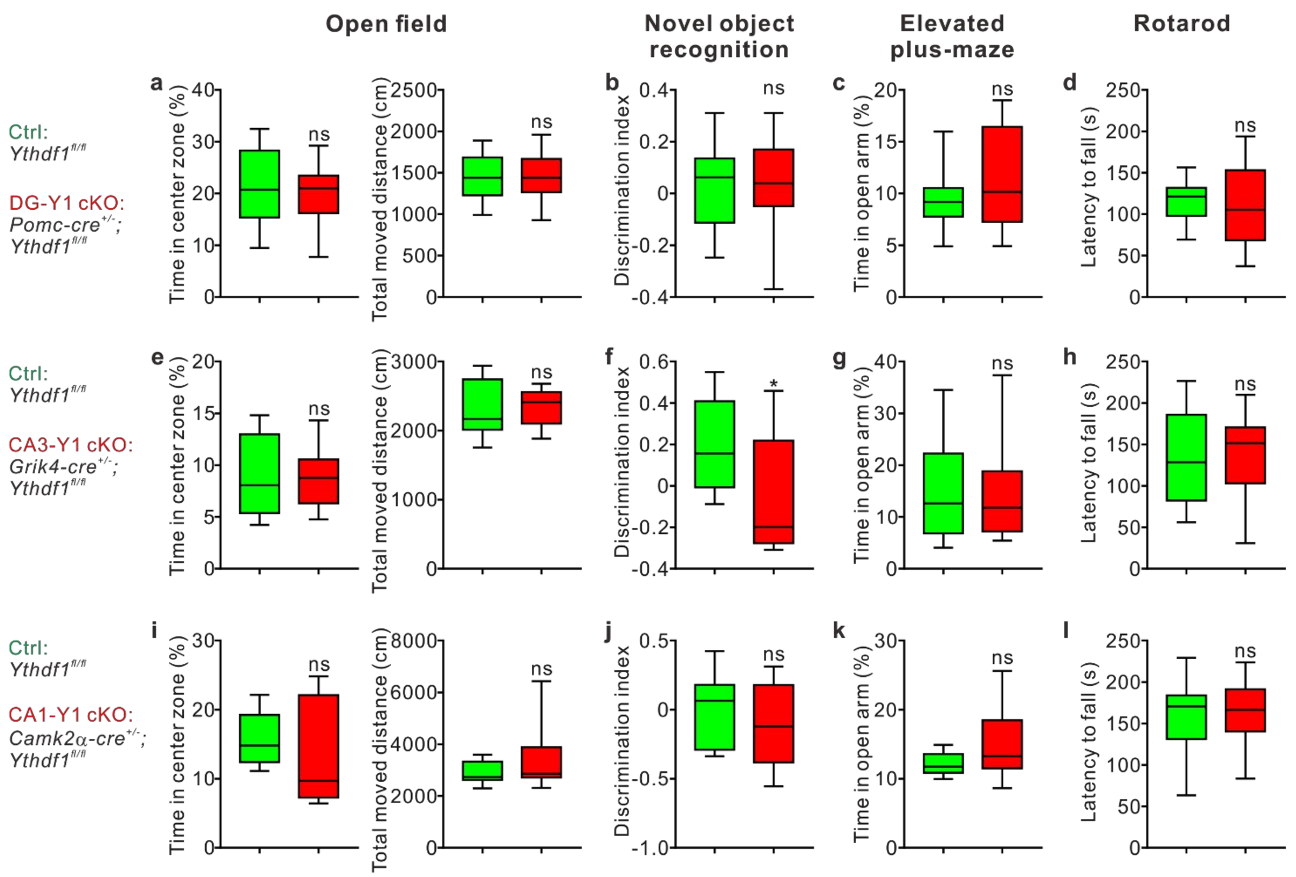


**Supplementary Fig. 5. DG-, CA3-, or CA1-specific *Ythdf1* conditional knockout mice are normal in other behavioral tests.**

DG-, CA3-, and CA1-specific *Ythdf1* cKO are designated as DG-Y1 cKO (a-d), CA3-Y1 cKO (e-h) and CA1-Y1 cKO (i-l), respectively. **a, e, i,** Exploratory behavior of these cKO mice and their controls was measured in the open field test. Mice with higher levels of exploration spend more time in the center zone. **b, f, j,** Short-term memory of these cKO mice and their controls was measured with the novel object recognition test. The amount of time that mice take to explore the known and unknown objects is used to calculate the discrimination index. **c, g, k,** Anxiety level of these cKO mice and their controls was measured in the elevated plus-maze test. The duration that mice spend in the open arm reflects their anti-anxiety behavior. **d, h, l,** Motor coordination of these cKO mice and their controls was measured with the rotarod test. Better motor coordination prevents mice from falling, resulting in longer latency to fall. No difference between cKO mice and their controls was observed in the above tests except that CA3-Y1 *cKO* mice showed impaired short-term memory compared with control mice. All data are represented as box and whisker plots. For DG-Y1 cKO, *n* = 13 mice for each genotype in a-d; for CA3-Y1 cKO, *n* = 14 mice for Ctrl and *n* = 16 mice for CA3-Y1 cKO in e-h, *$P$ = 0.016 in f; for CA1-Y1 cKO, *n* = 10 mice for each genotype in i-l; ns, not significant; all by unpaired Student's t-test.

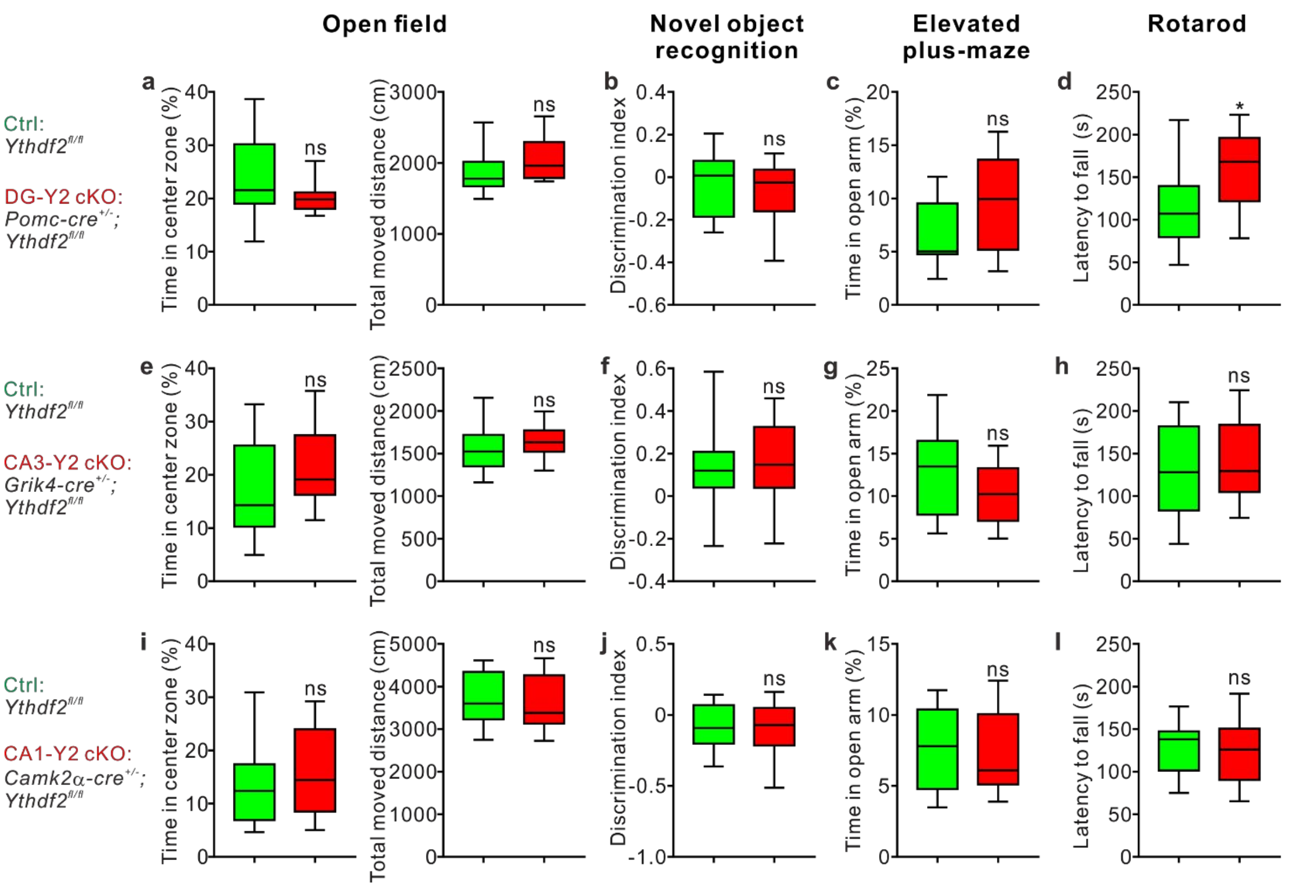


**Supplementary Fig. 6. DG-, CA3-, or CA1-specific *Ythdf2* conditional knockout mice are largely normal in other behavioral tests.**

DG-, CA3-, and CA1-specific *Ythdf2* cKO are designated as DG-Y2 cKO (a-d), CA3-Y2 cKO (e-h), and CA1-Y2 cKO (i-l), respectively. **a, e, i,** Exploratory behavior of these cKO mice and their controls was measured in the open field test. **b, f, j,** Short-term memory of these cKO mice and their controls was measured with the novel object recognition test. **c, g, k,** Anxiety level of these cKO mice and their controls was measured in the elevated plus-maze test. **d, h, l,** Motor coordination of these cKO mice and their controls was measured with the rotarod test. No difference was observed between cKO mice and their controls in the above tests except that DG-Y2 *cKO* mice showed better motor coordination than control mice. All data are represented as box and whisker plots. For DG-Y2 cKO, $n$ = 14 mice for Ctrl and $n$ = 10 mice for DG-Y2 cKO in a-d, *$P$ = 0.036 in d; for CA3-Y2 cKO, $n$ = 16 mice for Ctrl and $n$ = 14 mice for CA3-Y2 cKO in e-h; for CA1-Y2 cKO, $n$ = 12 mice for Ctrl and $n$ = 16 mice for CA1-Y2 cKO in i-l; ns, not significant; all by unpaired Student's t-test.

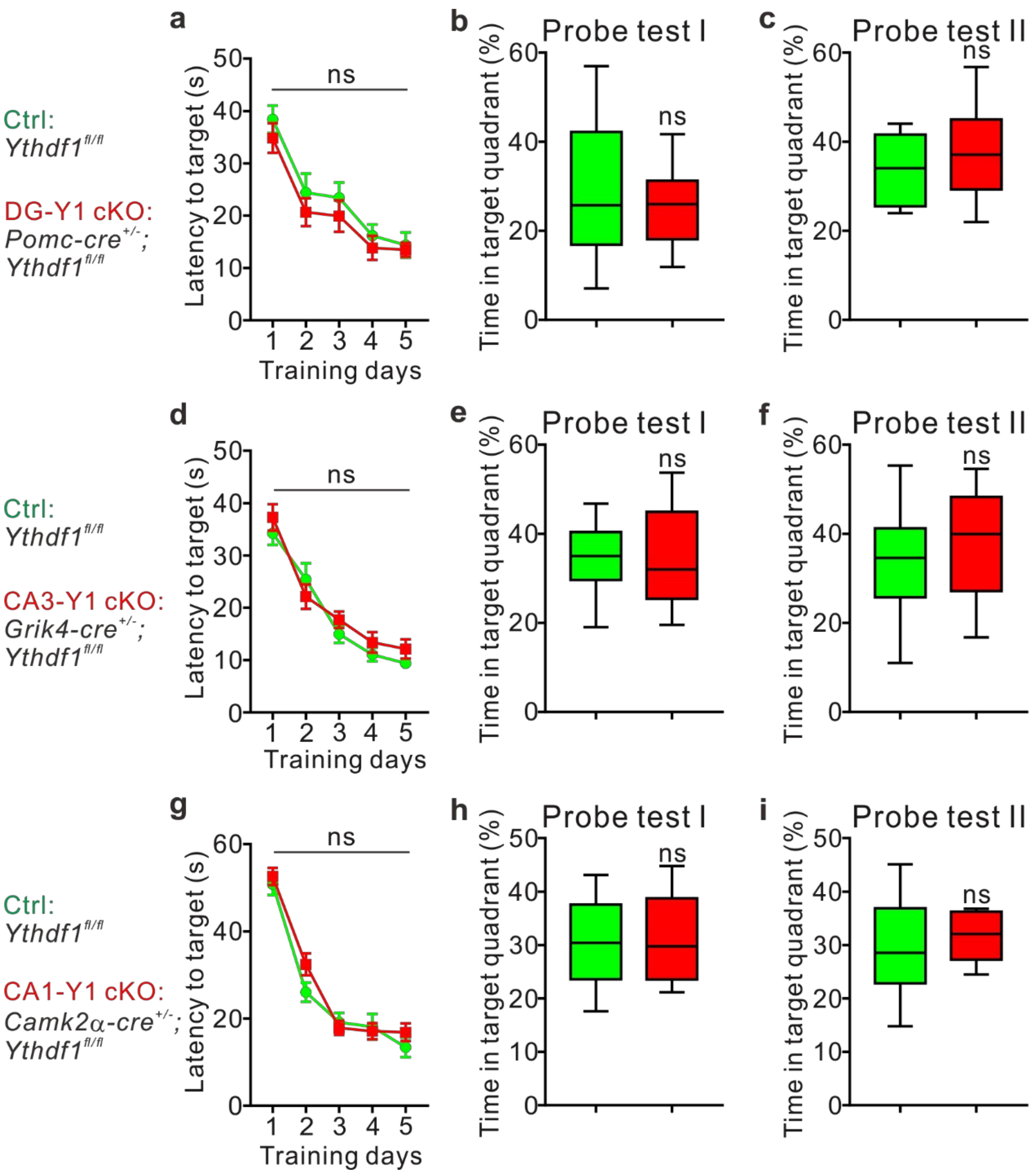


**Supplementary Fig. 7. Learning and memory are not disturbed in *Ythdf1* conditional knockout mice.** DG-, CA3-, and CA1-specific *Ythdf1* cKO mice are designated as DG-Y1 cKO (a-c), CA3-Y1 cKO (d-f), and CA1-Y1 cKO (g-i). The performance of these cKO mice showed no difference from their controls in the Morris water maze test by quantification of learning curves (a, d, g), Probe test I on day 3 (b, e, h), and Probe test II on day 6 (c, f, i). Quantifications of Probe tests I-II are represented as box and whisker plots, while the learning curves are shown in dot plots (mean ± SEM). For DG-Y1 cKO, *n* = 13 mice for each genotype in a-c; for CA3-Y1 cKO, *n* = 14 mice for Ctrl and *n* = 16 mice for CA3-Y1 cKO in d-f; for CA1-Y1 cKO, *n* = 10 mice for each genotype in g-i; ns, not significant; all box and whisker plots by unpaired Student's t-test; group difference in dot plots by two-way ANOVA with Tukey's post hoc test.

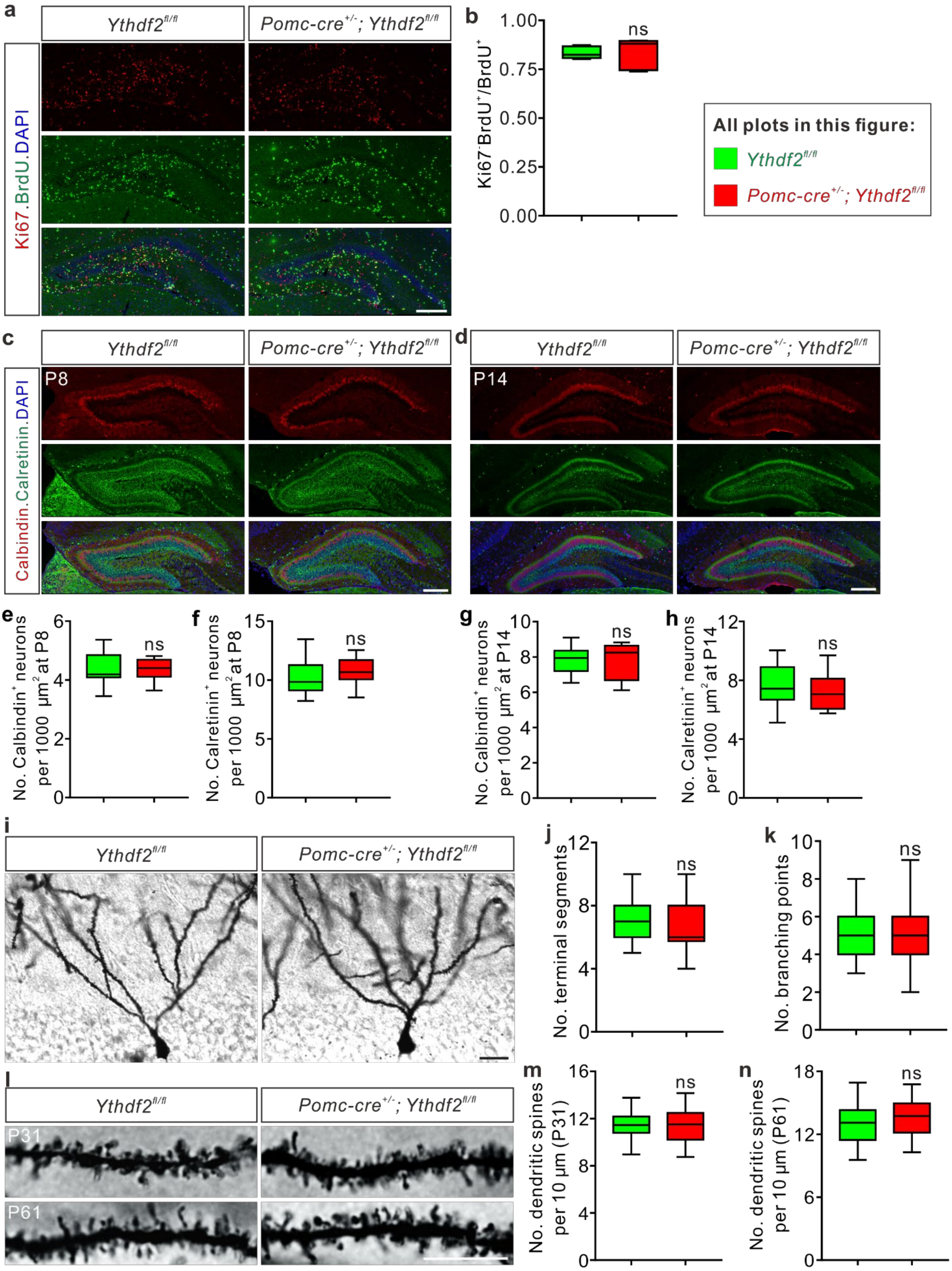


**Supplementary Fig. 8. Neurogenesis or dendrite development of DG granule cells are not affected in the DG-specific *Ythdf2* knockout mice.**

**a, b,** Representative images of Ki67 and BrdU labelling of P8 DG. Quantification of ratios of Ki67$^-$ cells in all

$BrdU^+$ cells in the DG area, which represent the new-born cells exiting cell cycle, showed no difference between DG-specific *Ythdf2* cKO and control mouse pups. **c, d,** Co-staining of Calretinin (green), a marker of immature granule cells (GCs), and Calbindin (red), a marker of mature GCs, in P8 (c) and P14 (d) DG showed no difference between DG-specific *Ythdf2* cKO and control mouse pups. **e-h,** Quantification of mature $Calbindin^+$ (e, g) and immature $Calretinin^+$ (f, h) granule cell numbers in DG of *Ythdf2* cKO and control mice at P8 (e, f) and P14 (g, h). **i-k,** Golgi staining revealed comparable morphology of GC dendrites in DG-specific *Ythdf2* cKO and their controls at P31 (i). Quantification showed the numbers of terminal segments and branching (secondary and tertiary branches) points of dendrites, which were not changed (j, k). **l-n,** Representative Golgi staining images showing the dendritic spines of GCs in DG-specific *Ythdf2* cKO and their controls at P31 and P61 (l). Quantification showed that *Ythdf2* cKO mice had comparable dendritic spine densities with control mice at P31 (m) and P61 (n). All data are represented as box and whisker plots: in b, *n* = 5 sections for $Ythdf2^{fl/fl}$, *n* = 7 sections for $Pomc\text{-}cre^{+/-}$; $Ythdf2^{fl/fl}$, *P* = 0.86; in d and e, *n* = 14 confocal fields for $Ythdf2^{fl/fl}$, *n* = 13 confocal fields for $Pomc\text{-}cre^{+/-}$; $Ythdf2^{fl/fl}$, *P* = 0.90 (d), *P* = 0.30 (e); in g and h, *n* = 12 confocal fields for $Ythdf2^{fl/fl}$, *n* = 12 confocal fields for $Pomc\text{-}cre^{+/-}$; $Ythdf2^{fl/fl}$, *P* = 0.95 (g), *P* = 0.53 (h); in j and j, *n* = 31 neurons for $Ythdf2^{fl/fl}$, *n* = 30 neurons for $Pomc\text{-}cre^{+/-}$; $Ythdf2^{fl/fl}$, *P* = 0.30 (j), *P* = 0.60 (k); in m, *n* = 54 dendrites for each genotype, *P* = 0.78; in m, *n* = 50 dendrites for each genotype, *P* = 0.074; ns, not significant; all by unpaired Student's t test. At least 3 mice were analyzed for each genotype in each experiment. Scale bars, 200 μm (a, c, d), 50 μm (i), 10 μm (l).

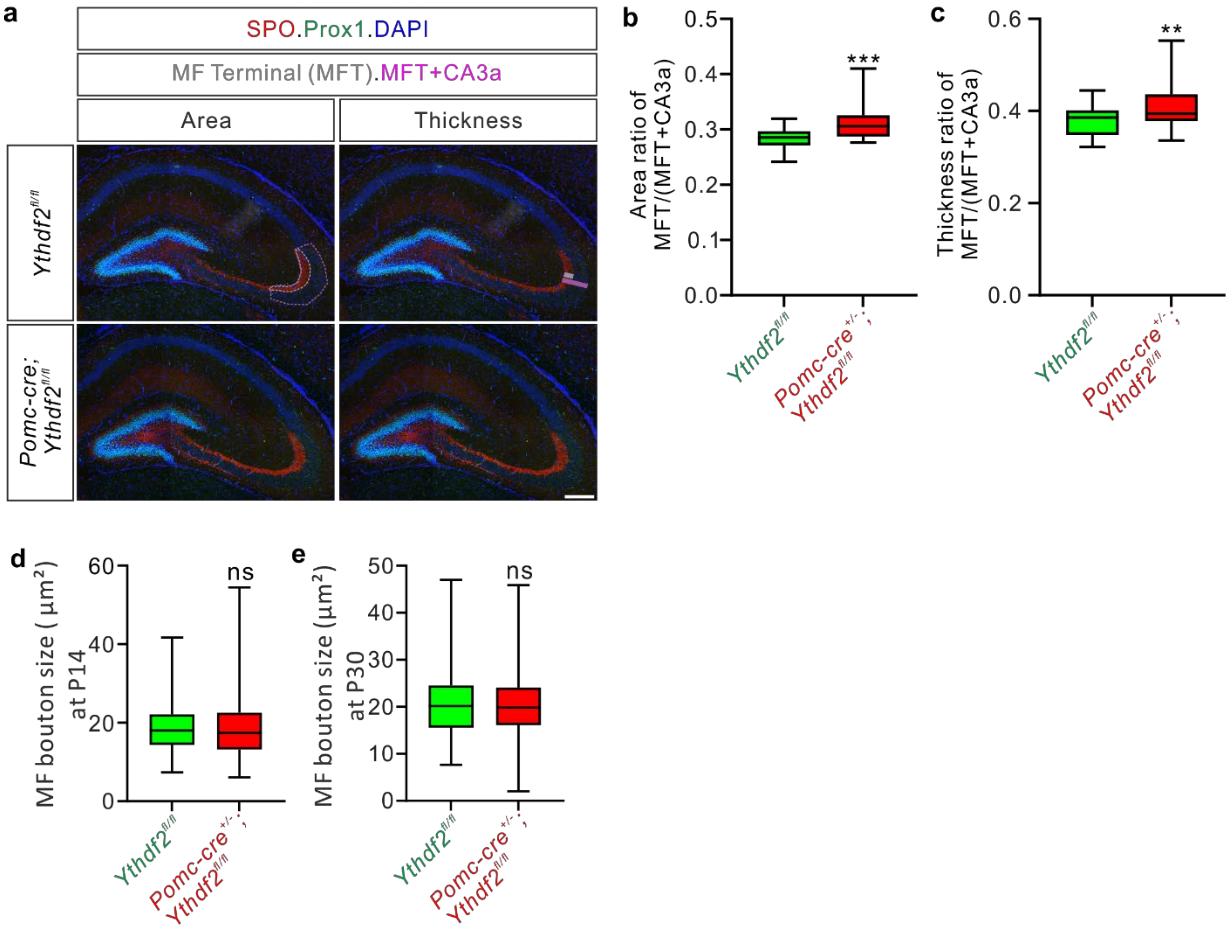


**Supplementary Fig. 9. Mossy fiber (MF) terminal is larger and MF bouton size is not affected in DG-specific *Ythdf2* knockout mice.**

**a-c,** MF terminal (MFT) was visualized by immunostaining of SPO, an MFT marker, at P8 (a). The grey and magenta dotted boundaries indicate the MFT area and the “MFT+CA3a” area, respectively, while the grey and magenta bars measure the thickness of MFT and “MFT+CA3a”, respectively. Quantifications are shown in b and c. **d, e,** DiI labeling was used to visualize mossy fiber boutons in the stratum lucidum, which showed no difference in bouton sizes between *Ythdf2*$^{fl/fl}$ and *Pomc-cre*$^{+/-}$*; Ythdf2*$^{fl/fl}$ mice at P14 (d) and P30 (e). All quantifications are represented as box and whisker plots: in b and c, *n* = 32 sections for *Ythdf2*$^{fl/fl}$*; n* = 29 sections for *Pomc-cre*$^{+/-}$*; Ythdf2*$^{fl/fl}$, ****P* = 1.80E-4 (b), ***P* = 0.0029 (c); in d, *n* = 217 boutons for *Ythdf2*$^{fl/fl}$, *n* = 207 boutons for *Pomc-cre*$^{+/-}$*; Ythdf2*$^{fl/fl}$, *P* = 0.50; in e, *n* = 261 boutons for *Ythdf2*$^{fl/fl}$, *n* = 269 boutons for *Pomc-cre*$^{+/-}$*; Ythdf2*$^{fl/fl}$, *P* = 0.91; ns, not significant; all by unpaired Student's t-test. At least 3 mice were analyzed for each genotype in each experiment. Scale bar, 200 μm (a).

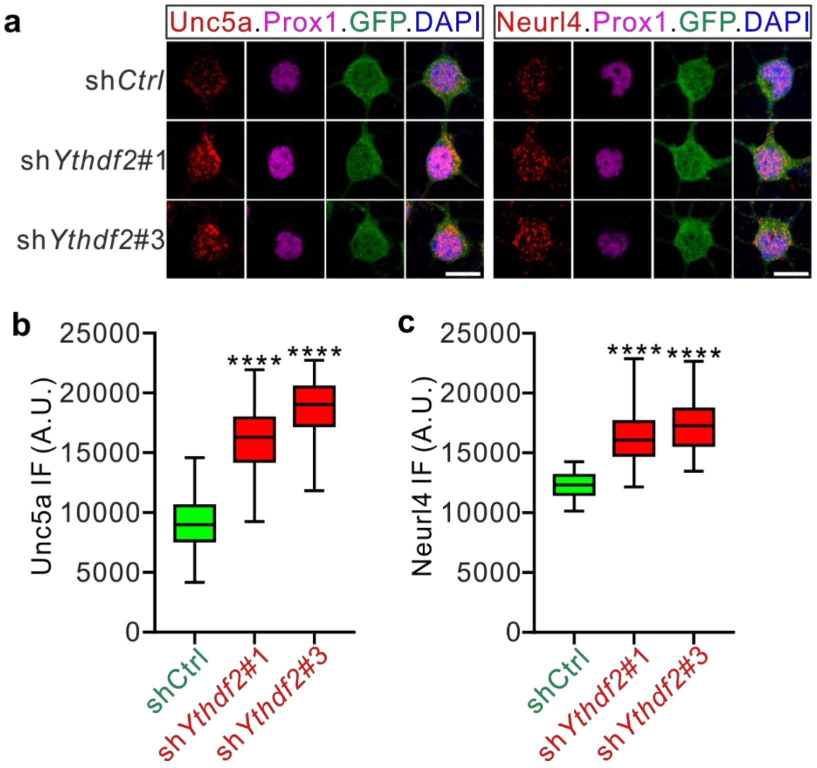


**Supplementary Fig. 10. Upregulation of Unc5a and Neurl4 protein levels after KD of YTHDF2 in cultured GC neurons.**

GC neurons dissected from P6-8 DG were cultured and infected with lentiviral sh*Ythdf2*, followed by immunostaining with Unc5a and Neurl4 antibodies (a). The IF signals were quantified (b, c). Quantifications are represented as box and whisker plots: in b, shCtrl ($n$ = 70 neurons) vs sh*Ythdf2*#1 ($n$ = 68 neurons), ****$P$ = 1.01E-40; shCtrl vs sh*Ythdf2*#3 ($n$ = 70 neurons), ****$P$ = 5.16E-59; in c, $n$ = 51 neurons for each group, ****$P$ = 2.11E-19 for shCtrl vs sh*Ythdf2*#1, ****$P$ = 8.21E-26 for shCtrl vs sh*Ythdf2*#3; by one-way ANOVA followed by Tukey's post hoc test. Scale bars, 20 μm (a).
